# An ultralight head-mounted camera system integrates detailed behavioral monitoring with multichannel electrophysiology in freely moving mice

**DOI:** 10.1101/294397

**Authors:** Arne F. Meyer, Jasper Poort, John O’Keefe, Maneesh Sahani, Jennifer F. Linden

**Author notes:** These authors contributed equally (Arne F. Meyer, Jasper Poort).

## Abstract

Breakthroughs in understanding the neural basis of natural behavior require neural recording and intervention to be paired with high-fidelity multimodal behavioral monitoring. An extensive genetic toolkit for neural circuit dissection, and well-developed neural recording technology, make the mouse a powerful model organism for systems neuroscience. However, methods for high-bandwidth acquisition of behavioral signals in mice remain limited to fixed-position cameras and other off-animal devices, complicating the monitoring of animals freely engaged in natural behaviors. Here, we report the development of an ultralight head-mounted camera system combined with head-movement sensors to simultaneously monitor eye position, pupil dilation, whisking, and pinna movements along with head motion in unrestrained, freely behaving mice. The power of the combined technology is demonstrated by observations linking eye position to head orientation; whisking to non-tactile stimulation; and, in electrophysiological experiments, visual cortical activity to volitional head movements.

## Introduction

A fundamental goal of neuroscience is to understand how neural circuits integrate a wide range of inputs to produce flexible and adaptive behaviors in natural settings. To approach this goal in its most general form, it will be essential to monitor and manipulate both neural activity and behavioural variables while animals interact naturally with their environments. The availability of genetic tools to dissect neural circuitry (Luo et al., 2008) and to construct models of human disease (Gatz and Ittner, 2008; Nestler and Hyman, 2010; Chesselet and Richter, 2011) has driven the emergence of the mouse as a key model organism in systems neuroscience (Carandini and Churchland, 2013). An increasingly wide array of technologies are available to measure and manipulate neural activity in mice (Voigts et al., 2008; Luo et al., 2008; Kim et al., 2017; Jun et al., 2017). However, detailed monitoring of behavior, especially in freely moving animals, remains a major challenge (Krakauer et al., 2017; Juavinett et al., 2018). To address this challenge, we developed an ultralight head-mounted camera system to measure eye position, pupil dilation, whisking, pinna movements and other behavioral signals in freely moving mice, which we combined with head-movement monitoring and multichannel electrophysiology.

Despite the longstanding ability to record neural activity in unrestrained rodents (e.g., O’Keefe and Dostrovsky, 1971), many current studies of the neural basis of behaviour have relied on awake but head-restrained animals (Carandini and Churchland, 2013; Juavinett et al., 2018). Head fixation enables tight control of sensory inputs, facilitates intracranial recording or imaging, and simplifies experimental manipulations that would be diffi cult in freely moving animals. However, results obtained in head-restrained animals may not generalize to more natural sensory and behavioral conditions (Tatler and Land, 2011; Felsen and Dan, 2005). For example, the change in vestibular inputs following head fixation may have widespread effects throughout the brain (Rancz et al., 2015), and it is debated whether spatial navigation by head-fixed animals in virtual reality environments is comparable to spatial navigation in freely moving animals (Dombeck et al., 2007; Chen et al., 2013; Domnisoru et al., 2013; Schmidt-Hieber and Häusser, 2013; Aghajan et al., 2015; Minderer et al., 2016). While the level of experimental control and the availability of techniques for monitoring neural activity are more limited in studies of freely moving animals, such investigations have provided important insights into brain function during behavior that might not have been obtained in more constrained experimental settings, for instance revealing cells that represent an animal’s spatial location and head direction (O’Keefe and Dostrovsky, 1971; Taube et al., 1990; Fyhn et al., 2004).

Detailed behavioral measurement in freely moving mice remains a major challenge because of the animal’s small size (average adult weight only *∼*20-25 grams; www.jax.org). Externally mounted video cameras have been used to track aspects of gross locomotor behavior including gait (Machado et al., 2015) and posture (Hong et al., 2015; Wiltschko et al., 2015), and (in semi-stationary mice and when permitted by the camera angle) whisking (Voigts et al., 2008; Roy et al., 2011; Nashaat et al., 2017) and head and eye movements (Kretschmer et al., 2015, 2017). However, the perspective of the external camera limits the potential for continuous measurement of whisking, pupil diameter or eye position in actively exploring mice (although Payne and Raymond, 2017, have successfully monitored horizontal eye movements using a magnetic field approach).

The new miniaturized head-mounted tracking system reported here makes it possible to continuously monitor multiple behavioral variables, such as eye and pinna movements, whisking, eating and licking, together with head movements, in combination with chronic neural recording from unrestrained mice. A recent study developed a head-mounted eye tracking system for the rat (Wallace et al., 2013). However, given the comparatively small size of the mouse, we required a system with greatly reduced weight and footprint. Moreover, the method used in rats relied on detection of reference points recorded by multiple video cameras and additional head-mounted LEDs to track orientation and movement of the head. Instead, we used small lightweight inertial sensors to track the orientation and movements of the head (Mizell, 2003; Pasquet et al., 2016), simplifying the process of relating these variables to the camera outputs even under demanding natural conditions.

The system generates stable video output, leaves mouse behavior largely unchanged, and does not affect the quality of concomitant neural recordings. We demonstrate the potential of the system in a series of experiments in freely moving mice. First, we show that variables such as whisking frequency and pupil size vary systematically with behavioral state, and that these changes are correlated with neural activity, thereby generalizing results obtained in head-restrained mice to natural behaviors (Reimer et al., 2014; McGinley et al., 2015). Second, we demonstrate that a large fraction of variability in eye position in freely moving mice is explained by head movements, as has also been observed in rats (Wallace et al., 2013). We find systematic relationships between eye position and head orientation in freely moving mice, suggesting that mice stabilize their gaze with respect to the horizontal plane, even in the dark. Third, we demonstrate that neural activity in primary visual cortex (V1) is strongly modulated by head movements even in the absence of visual input. This effect does not depend on variability in eye movements and cannot be explained by whisking or locomotion. These results demonstrate how the new camera system can lead to novel insight into the interactions between different behaviors and their relation with neural activity.

## Results

### A miniature head-mounted camera system for freely moving mice

The ultra-lightweight head-mounted camera system (Figure 1A) consisted of a miniature CMOS image sensor with integrated video data cable; a custom 3D-printed holder for the image sensor; an infrared (IR) LED illumination source; and an IR mirror on a custom lightweight extension arm. The mirror reflected only IR light and allowed visible light to pass through, so it was visually transparent to the mouse (Peirson et al., 2017). The weight of the camera system including the image sensor was approximately 1.3 grams (see Figure S1, Table S1, and Methods for the list of parts). We wrote custom software (see Methods) to synchronize video and neural data and to integrate video recordings with open-source systems for neural data acquisition (http://www.open-ephys.org). The camera system recorded video frames with image resolution 640×480 pixels at frame rates of up to 90 Hz (Figure S2); thus, video images could be aligned to neural data with a temporal precision of 11.1 ms.

**Figure 1:**
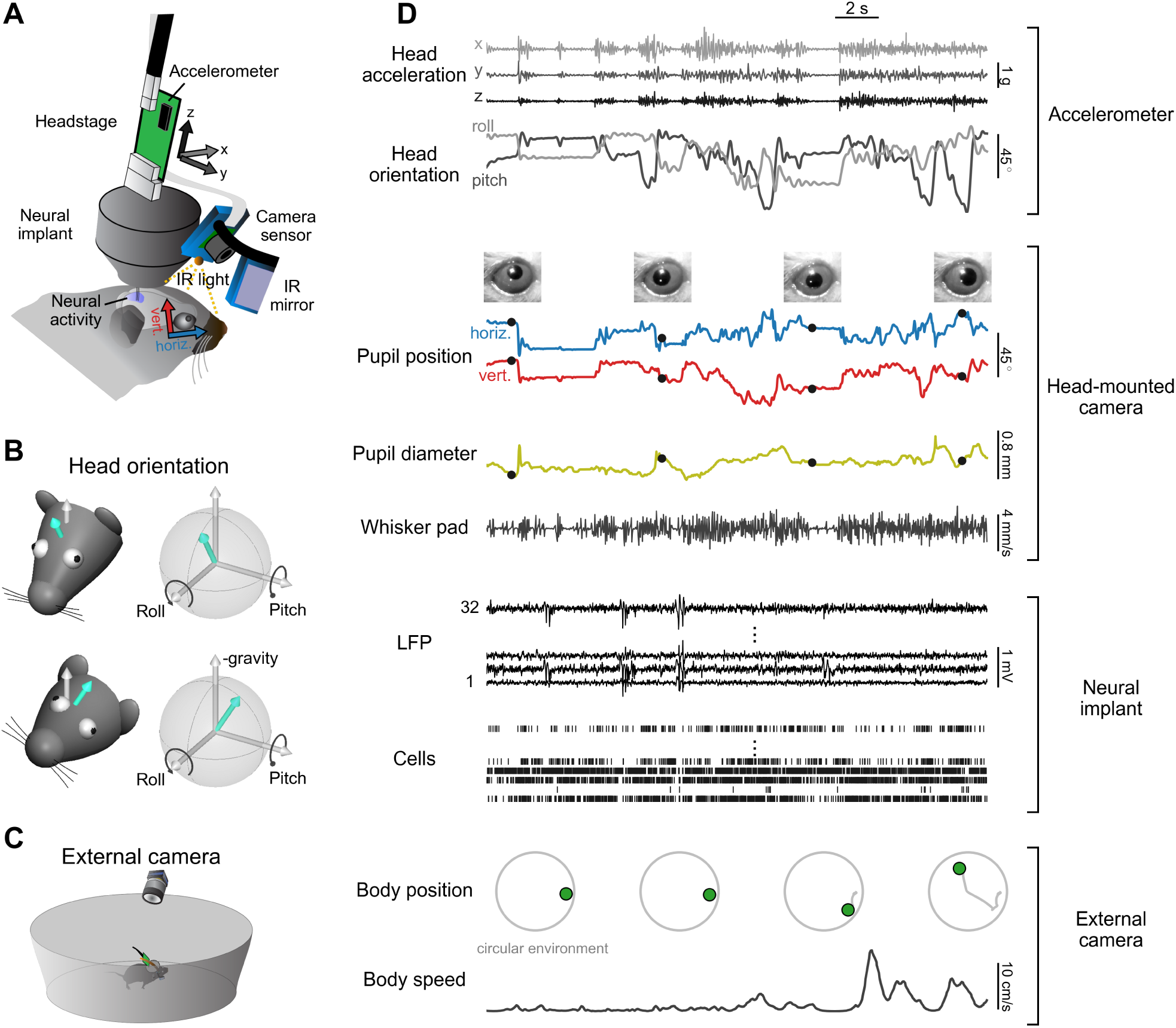
Simultaneous measurement of multiple behavioral variables and neural activity in a freely-moving mouse. (A) Neural activity recorded with a chronic tetrode implant; video data simultaneously recorded using a miniature CMOS image sensor and an infrared (IR) mirror mounted on the implant with a custom lightweight holder. An IR light source on the camera holder illuminates the region of interest, which is imaged via the IR mirror. The mirror reflects only IR light, allowing visible light to pass through so the animal’s vision is not obstructed. Head motion and orientation are measured using an accelerometer integrated into the neural recording headstage. (B) Extraction of pitch and roll from low-pass filtered accelerometer signals. White arrow indicates direction opposite to gravity component. Turquoise arrow indicates orientation of vertical (ventral-dorsal) head axis. (C) A mouse freely explores its environment while wearing the head-mounted camera system. Absolute position is measured using external cameras. (D) Example traces of simultaneously recorded behavioral and neural data. Pictures of eye position in third row were acquired at times of dots on pupil position traces in the fourth row.

The camera system was attached during each recording session to a miniature connector built into a chronically implanted custom tetrode drive with 8-16 individually movable tetrodes (based on an existing implant design, Voigts et al., 2013). Power to the IR LED was provided through the digital neural recording head-stage, which was also attached to the implant for each recording session. The headstage board included an integrated 3-axis accelerometer to measure the movement and orientation of the animal’s head (Pasquet et al., 2016) (Figure 1B and Methods; see Figure 7 and Methods for measurement of rotational movements). The mouse freely explored a small circular environment, while body position was monitored using an external camera (Figure 1C). The combined system allowed the simultaneous measurement of pupil position, pupil dilation, whisker pad movement, head movement, head orientation, body position and body speed together with neural activity in freely moving mice (Figure 1D).

### Operation of camera does not impair neural recording quality

We first checked that the camera system did not interfere with electrophysiological recordings, by comparing neural recordings with camera powered and operating or switched off. Figure 2A,B shows signals from an example tetrode channel. The isolation of action-potential spikes appeared unchanged with the camera on or off, and the projection of spike waveforms into a noise-whitened robust PCA (NWrPC) space (Sahani, 1999) was similar in both conditions (insets in Figure 2A,B) as was the power spectrum of the broadband signal (Figure 2C). We quantified spectral differences between conditions across recordings by computing the power ratio between counter-balanced camera-on and camera-off recording segments obtained during each recording session. Across the frequency range of neural signals (2 to 6000 Hz), differences in log-power ratios were close to 0 dB, and within one standard deviation of the within-condition (camera-off) variability (Figure 2D-F). We also compared the signal-to-noise ratio (SNR) for spike waveforms recorded from the same cells with the camera on or off (Figure 2G and Methods), and found no significant difference (Wilcoxon signed-rank test, P =0.19). Thus the operation of the head-mounted camera had no discernable impact on the signal quality of electrophysiological recordings.

**Figure 2:**
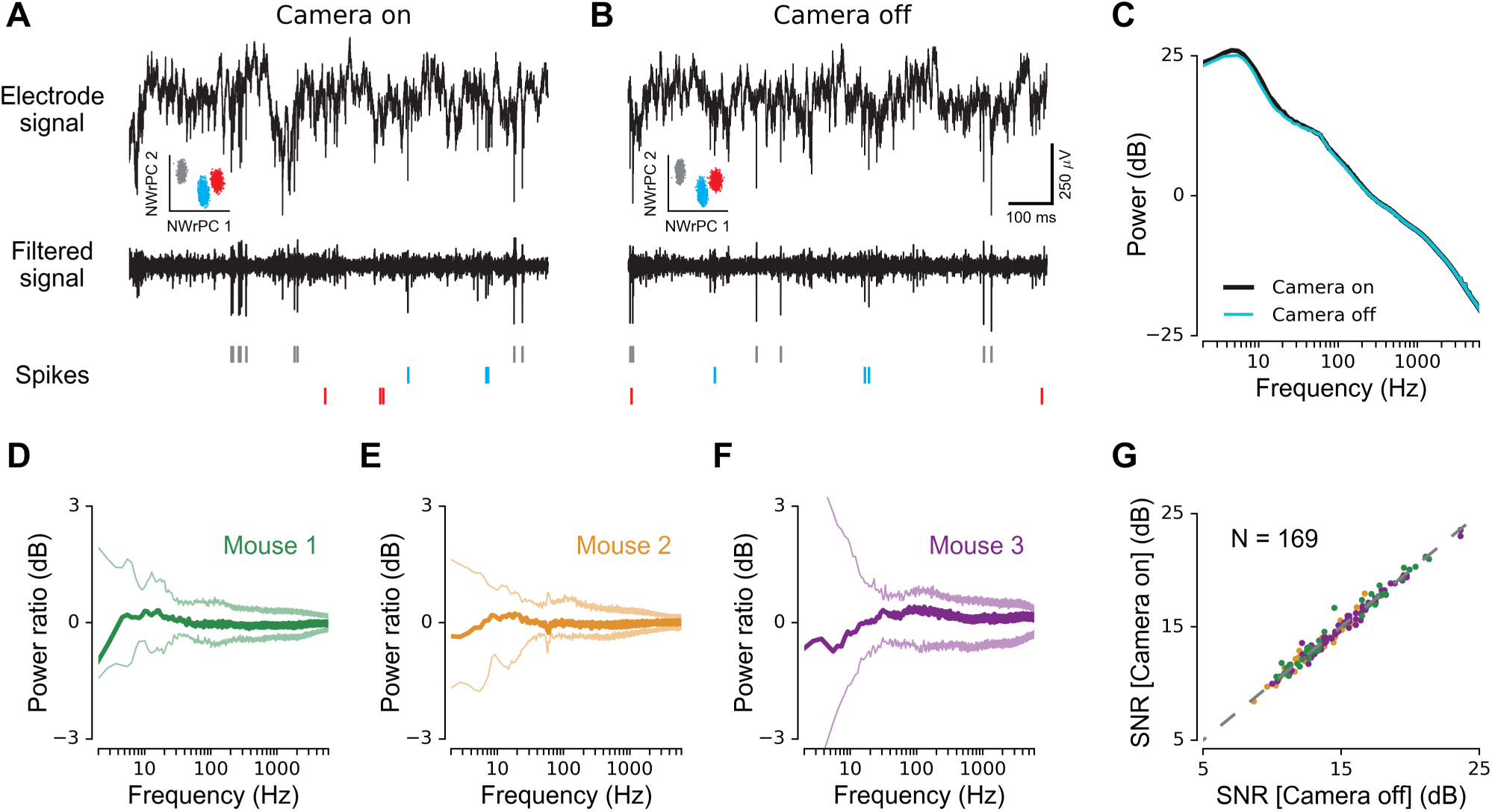
Neural recording quality with head-mounted camera. (A) Broadband electrode signal (top), high-pass filtered signal (middle), and extracted spikes (bottom) with head-mounted camera activated ("camera-on" condition). Inset shows projections of spike waveforms for identified cells into the space defined by the first two noise-whitened robust principal components (NWrPC 1 and 2). (B) The same as in A but with head-mounted camera de-activated ("camera-off" condition). Scales apply to bottom and top traces in A and B. (C) Power density spectrum of broadband electrode signals for camera-on and -off conditions (10 minutes each) recorded in the same session. Note that the two lines are closely overlapping. (D-F) Mean log power ratio between broadband electrode signals in camera-on and -off conditions (10 minutes each condition per session) across recording sessions for three mice (mouse 1 & 2, *n* = 9; mouse 3, *n* = 19). Pale lines indicate standard deviation of log power ratios for recordings in camera-off condition alone. (G) Signal-to-noise ratio (SNR) between identified spike waveforms and noise in the high-pass filtered signal, for camera-on versus camera-off conditions (158 single-units, 11 multi-units).

### Camera images remain stable as the mouse moves

Next, we measured the stability of video recordings from the head-mounted camera. We identified a rigid part of the implant visible in the image frame as a reference (grey outline in inset image in Figure 3A) and used motion registration (Dubbs et al., 2016) to determine the x- and y-displacement of the image in each frame, relative to the average image position across frames. When displacements occurred, they were typically on the order of a single pixel (40 *µ*m; Figure 3A). The diameter of the mouse eye and pupil are approximately 3.4 mm (Sakatani and Isa, 2004) and 0.4 - 1.6 mm (McGinley et al., 2015), respectively. Thus, on average, camera image displacements in freely exploring mice were 1-2 orders of magnitude smaller than eye or pupil diameter. Moreover, average inter-frame image movement (i.e., change in 2D displacement between successive frames) was less than 4 *µ*m in mice freely exploring a circular environment, compared to less than 1 *µ*m in a control condition when the same animals were head-fixed on a cylindrical treadmill (Figure 3B and Methods).

**Figure 3:**
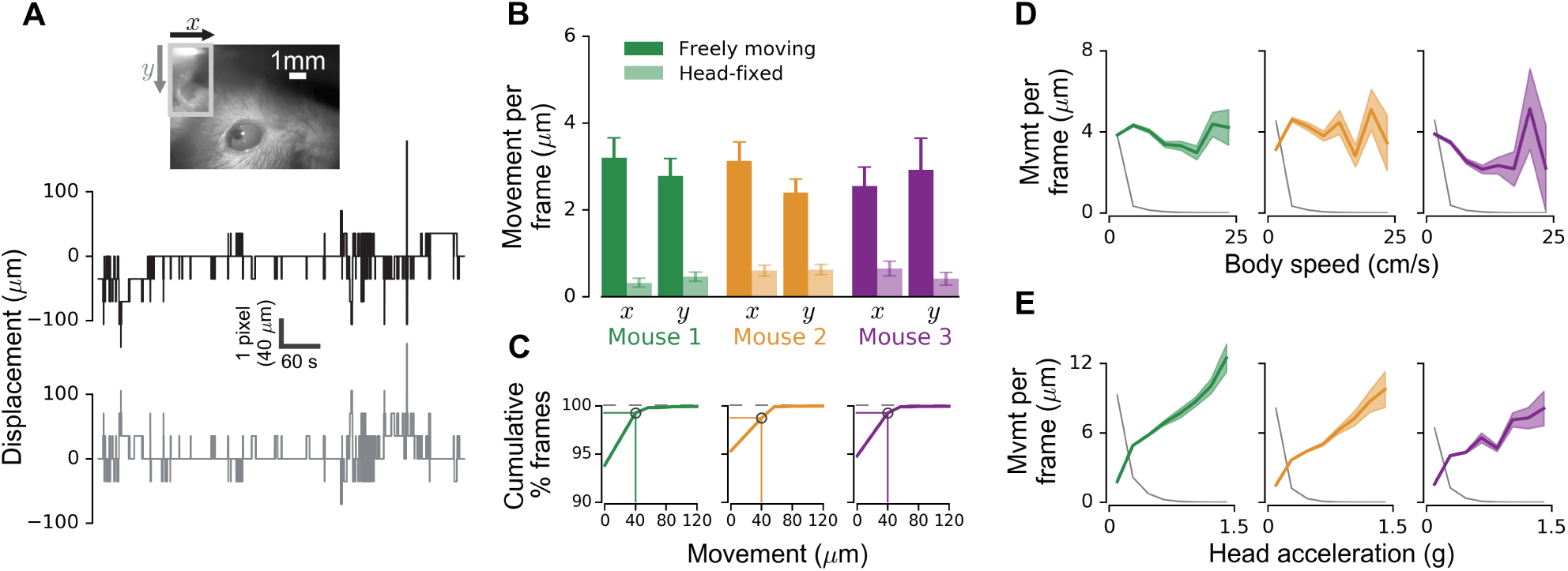
Image stability during movement. (A) Camera view of the left eye (top) with inset showing reference for image registration (gray rectangle). Traces below show example frame-by-frame displacements of camera image in x- (middle) and y- (bottom) directions. (B) Average 2D inter-frame image movement for three mice, recorded while animals were either freely exploring a circular environment or head-fixed on a cylindrical treadmill. Number of freely moving and head-fixed recordings (10 minutes each): mouse 1, *n* = 55 and 22; mouse 2, *n* = 35 and 29; mouse 3, *n* = 14 and 14, respectively. (C) Cumulative distribution of inter-frame image movements. Note that image movement is zero for nearly 95% of frames. (D-E) Average inter-frame image movement as a function of body speed (D) or head acceleration (E), for three mice. Thin grey lines indicate relative frequency of body speed (D) or head acceleration (E), respectively.

We also investigated the frequency with which image movements occurred in freely moving mice. Figure 3C shows the cumulative distribution of inter-frame image movements, after excluding frames in which the reference was occluded, e.g., during grooming (less than 0.6% of all frames, see Methods). In nearly 95% of analyzed frames, no image movement was observed. In 98-99% of frames, the maximal shift was one pixel (40 *µ*m; see marked points in Figure 3C).

Finally, we investigated whether image movement was related to mouse behavior. There was no evident relationship between average image movement per frame and body speed (Figure 3D). We also tested for a relationship with head acceleration (after removing the gravity component, see Figure 1D and Methods) and found an increase in image movement with stronger head accelerations, but these strong head movements were rare in all three mice (head acceleration magnitude less than 0.2 g for 95% of the recorded frames in all mice; Figure 3E). Moreover, even when mice made head movements with a magnitude of 1 g, the average image movement per frame did not exceed about 10 *µ*m.

We conclude that the head-mounted camera system produced stable video recordings, even when mice were grooming or actively exploring objects in complex and enriched environments (see Movie S2).

### Patterns of behavior are minimally disturbed by camera system

Previous work has shown that mice tolerate the tetrode implant with only minimal changes in natural behavior (Voigts et al., 2013). We wondered whether the additional weight of the head-mounted camera system might alter gross locomotor and exploratory behaviors in our animals. We analyzed the head-mounted accelerometer signals obtained from two implanted mice with and without the camera attached, during repeated sessions of free exploration across more than two months. We developed a semi-automatic state-segmentation algorithm to segment the recordings into four behaviors (active exploration, quiescence, grooming, eating) based on the short-term spectra of the accelerometer signals (see Methods, Figure 4B,C and Figure S4). We found that this approach more accurately matched human observer segmentation (with cross-validation) than approaches based on segmenting the time-domain accelerometer signals directly (Venkatraman et al., 2010; Dhawale et al., 2017) (Figure S4D,E). Cross-validated classifications of behavioral state using the spectra-based algorithm matched classifications by a human observer over 96% of the time both with and without the camera attached, with no significant difference in classification performance between the two conditions (Figure 4D, Fisher’s exact test, *P*=0.40; Figure S4A, *P*=0.13).

**Figure 4:**
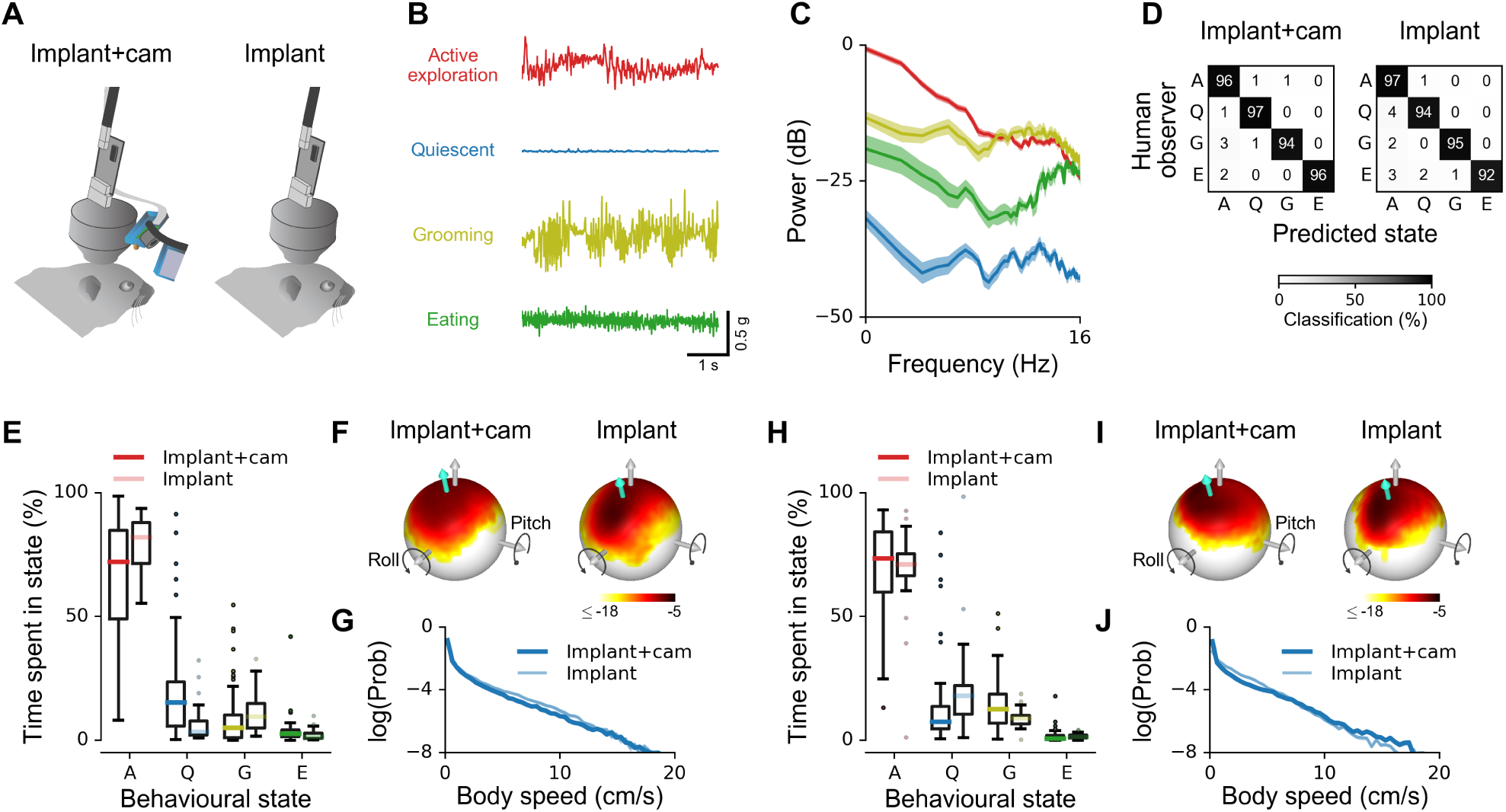
Impact of head-mounted camera on basic mouse behavior. (A) Recordings were performed with ("Implant+cam") and without ("Implant") head-mounted camera system. (B) Example accelerometer traces for one motion axis recorded in different behavioral states. (C) Power spectra of accelerometer signals shown in B, extracted from a 20-minute recording. The different behavioral states can be reliably discriminated based on the power spectra. Shaded areas indicate standard error. (D) Confusion matrix illustrating cross-validated classification performance of a semi-automatic state-segmentation algorithm based on head-mounted accelerometer signal spectra ("Predicted state"), compared to behavioral state classifications based on manual annotation of external video and other data ("Human observer"; see Methods). Left: mouse with implant and camera. Right: with implant only. (E) Distribution of proportions of time per session spent in different behavioral states for mouse 1. Dark and light colors of each hue indicate condition with and without camera, respectively. Number of sessions: implant+cam *n* = 21, implant alone *n* = 11. (F) Log-probability distribution of head orientation for the mouse in E, with implant and camera (left) and with implant alone (right). Gray arrow indicates direction opposite gravity; turquoise arrow indicates mean head orientation. (G) Log-probability distribution of measured body speed for mouse 1. (H-J) The same as in E-G for mouse 2. Number of sessions: implant+cam *n* = 18, implant alone *n* = 11.

The successful semi-automated segmentation of behavioral states allowed us to objectively compare mouse behavior with and without the camera. Behavioral patterns varied from day to day (Figure S4B,C), but both animals spent the majority of time in the active exploration state in most sessions (Figure 4E,H). The proportion of time spent in each behavioral state depended in part on session number relative to the first recording (Figure S4B,C). However, we found no statistically significant differences between implant+cam and implant alone conditions in the proportion of time spent in each state for either mouse (permutation test, *P*=0.07 for mouse 1, *P*=0.12 for mouse 2; see Methods for details). Each mouse divided its time similarly between the four behavioral states with and without the camera (Figure 4E,H).

Since the majority of time was spent in the active exploration state, we examined behavior in this state more closely, paying specific attention to head movements and body speed (Figure 4F,I). The addition of the camera produced a slight change in average head position (mouse 1: −7°pitch, +5°roll; mouse 2: −3°pitch, +5°roll), which was not statistically significant for either mouse (permutation tests; mouse 1: *P*=0.41 pitch, *P*=0.06 roll; mouse 2: *P*=0.92 pitch, *P*=0.37 roll). The camera also produced a small reduction in the standard deviation of head pitch, and a small increase in the standard deviation of head roll (mouse 1: +3°pitch SD, −4°roll SD; mouse 2: +4°pitch SD, −6°roll SD), each statistically significant in one of the two mice (permutation tests; mouse 1: *P*=0.04 pitch SD, *P*=0.12 roll SD; mouse 2: *P*=0.17 pitch SD, *P*=0.04 roll SD), and even here the differences were relatively small (−11% for pitch in mouse 1 and +30% for roll in mouse 2). Distributions of body speed during active exploration were unaffected by the camera (Figure 4G,J; permutation test, *P*=0.35 mouse 1, *P*=0.39 mouse 2; see Methods). We conclude that active exploratory head and body movements were minimally affected by the presence of the head-mounted camera.

### Pupil diameter and whisking correlate with behavioral and neural state in freely moving mice

We next explored the capacity of the combined implant and camera system to identify correlations between behavioral and neural variables. Figure 5A shows a 6-minute extract from a 40-minute recording session of several behavioral and neural variables which included active and quiescent states, as well as grooming and eating (see Movie S3 for a longer 10-minute segment).

**Figure 5:**
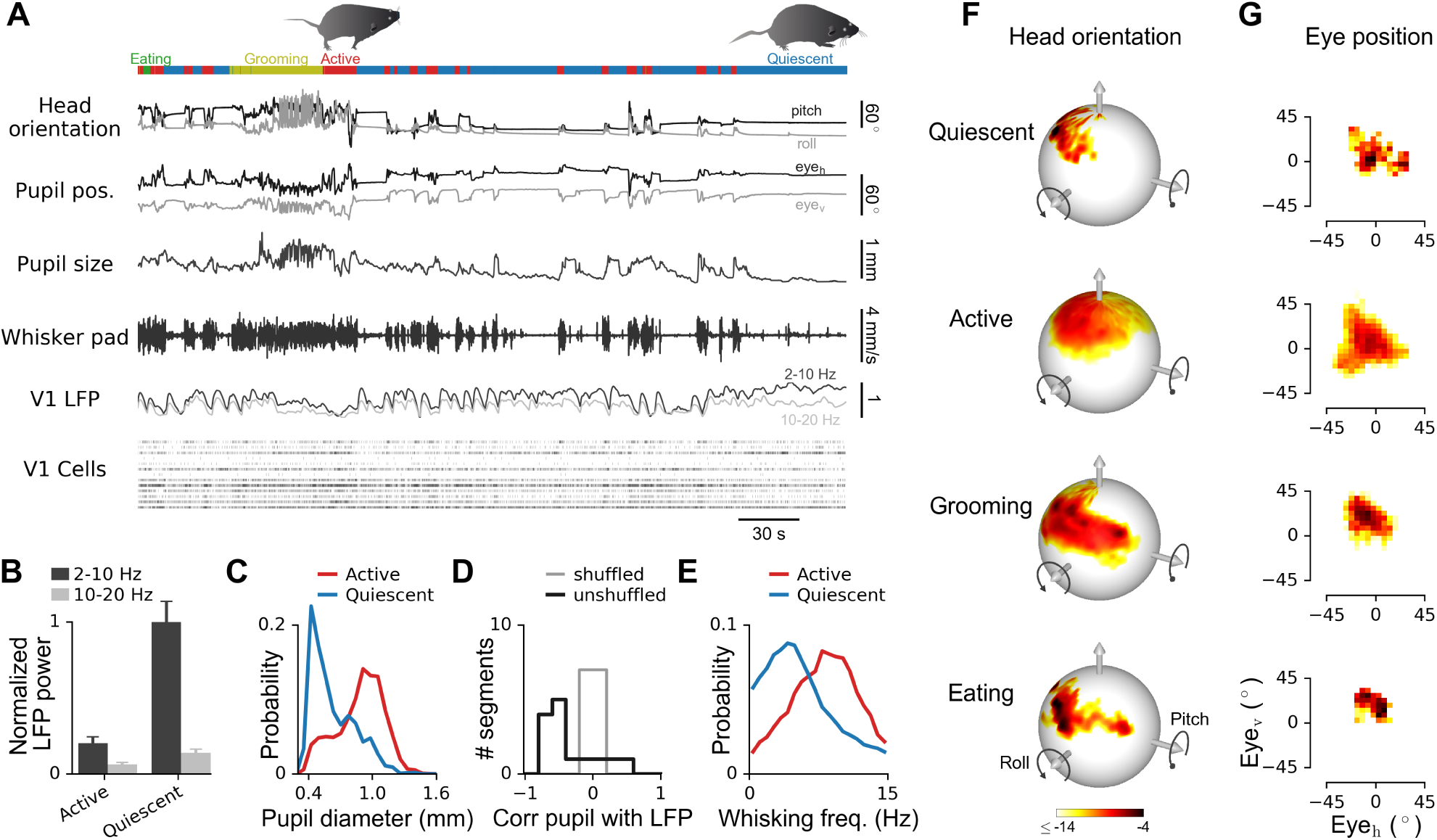
Continuous monitoring of behavioral and neural variables in freely moving mice. (A) Example traces of simultaneously measured behavioral and neural variables (5 minutes from a 40-minute recording). Colored rectangles above traces indicate behavioral states assigned by the behavioral segmentation algorithm. (B) Low (2-10 Hz) and high frequency (10-20 Hz) LFP power in V1 in active and quiescent states (mean ± SEM). LFP power normalized by low frequency power in quiescent state. (C) Distribution of pupil diameters in active and quiescent states. (D) Correlation coeffi cient between low-frequency (2-10 Hz) LFP power and pupil diameter during quiescent state. Only segments in which the head was still for at least 15 seconds were used for the analysis. (E) Distribution of whisker pad movement frequencies in active and quiescent states (30 Hz frame rates). (F) Log-probability distributions of head orientation in different behavioral states. (G) Log-probability distributions of simultaneously measured horizontal and vertical eye position in the same states. Same colorbar as in F.

Previous studies in head-restrained mice have demonstrated that local field potential (LFP) power in the low-frequency (2-10 Hz) band is significantly reduced in the active compared to the quiescent state in sensory cortex (Poulet and Petersen, 2008; McGinley et al., 2015). Moreover, in head-fixed animals, pupil diameter is inversely related to low-frequency LFP power, and increased during active behavior and reduced during quiescence (Reimer et al., 2014; McGinley et al., 2015). We found that these relations also hold in primary visual cortex (V1) in freely moving mice (Figure 5B-D). Normalized low-frequency LFP power was significantly lower in the active than quiescent state (Figure 5B; two-sample *t*-test, *P* <0.001), and the distribution of pupil diameters was shifted to larger values in the active state (Figure 5C; Wilcoxon rank-sum tests for difference in medians, *P* <0.001). Low-frequency LFP power and pupil diameter were not only inversely affected by changes between active and quiescent behavioral states, but also negatively correlated in simultaneous recordings within the same behavioral state. We analyzed correlations between LFP power and pupil diameter for quiescent recording segments during which the mouse kept its head in a constant position for at least 15 seconds, to minimize fluctuations in pupil diameter from changes in eye illumination (see also Methods). There was a strong negative correlation between pupil diameter and low-frequency LFP power in these recordings (Figure 5D; median correlation coeffi cient −0.44 versus 0 for shuffl ed data, Wilcoxon rank-sum test *P*=0.02).

Previous studies in head-restrained mice have also reported that the frequency of whisking is increased in the active compared to the quiescent behavioral state (Moore, 2004; Poulet and Petersen, 2008; Reimer et al., 2014). To examine whisking frequency in freely moving mice, we extracted whisker pad movements from the head-mounted camera images (see Methods and Figure S3 for details) and observed an increased frequency of whisker pad movements in the active state (Figure 5E; Wilcoxon rank-sum test for Wilcoxon rank-sum tests for difference in medians, *P* <0.001), confirming previous findings in head-restrained mice. We also discovered an aspect of whisking behavior that has not, to our knowledge, been reported previously in head-restrained mice: sounds that were presented when the mouse was immobile reliably evoked whisker pad movements that were comparable in magnitude to whisker pad movements observed during active exploration (Figure S5).

The head-mounted camera system also enabled measurement and analysis of head movements and head-movement-related behavior, which cannot be studied in head-restrained animals. We measured the distributions of head orientation (Figure 5F) and eye position (Figure 5G) in four behavioral states (quiescent, active, grooming and eating), by segmentation of behavioral data from continuous 40-minute recording sessions (see Figure 4A-D). The distributions of both head orientation and eye position had wider spreads during active exploration than during quiescence (Figure 5F,G; permutation test, *P* <0.001 for head pitch/roll and horizontal/vertical eye positions; see Methods). More specifically, the distributions in the quiescent state appeared to be dominated by particular combinations of head orientation and eye position that the mouse preferred at rest. In contrast, there was a different pattern during grooming: distinct modes of head orientation (which appeared to correspond to different grooming movements, e.g. forepaws over the nose and muzzle, strokes with the hindleg), combined with the same modal eye position (Figure 5F,G). Similarly, eye position remained relatively constant during eating, despite changes in head orientation. These observations indicate that head-eye coordination differs between behavioral states; eye-movement patterns are more restricted relative to head orientation during grooming and eating than during active exploration.

These results demonstrate that the head-mounted camera system enables detailed characterization of the relationship between multiple behavioural variables and neural activity in freely behaving mice. In addition, it can help to reveal subtle aspects of natural behavior, such as sound-evoked whisking movements and differences in head-eye coordination between behavioral states.

### Eye position depends on head orientation in freely moving mice

We wondered if the broader distribution of eye positions in actively exploring mice (Figure 5G and Figure 6A, top) compared to quiescent mice (Figure 5G) or head-restrained mice moving on a cylindrical treadmill (Figure 6A, bottom, Movie S5) was related to the larger range of head orientations during active exploration (Figure 5F). Previous results in head-restrained mice (Andreescu et al., 2005; Oommen and Stahl, 2008) and freely moving rats (Wallace et al., 2013) have suggested that average eye position varies systematically with orientation of the head. Indeed, in head-restrained, passively rotated mice, eye position varies systematically with head pitch and roll (Oommen and Stahl, 2008). Therefore, we used head-mounted accelerometers to measure head orientation (pitch and roll) (Figure 6B,C and Methods).

**Figure 6:**
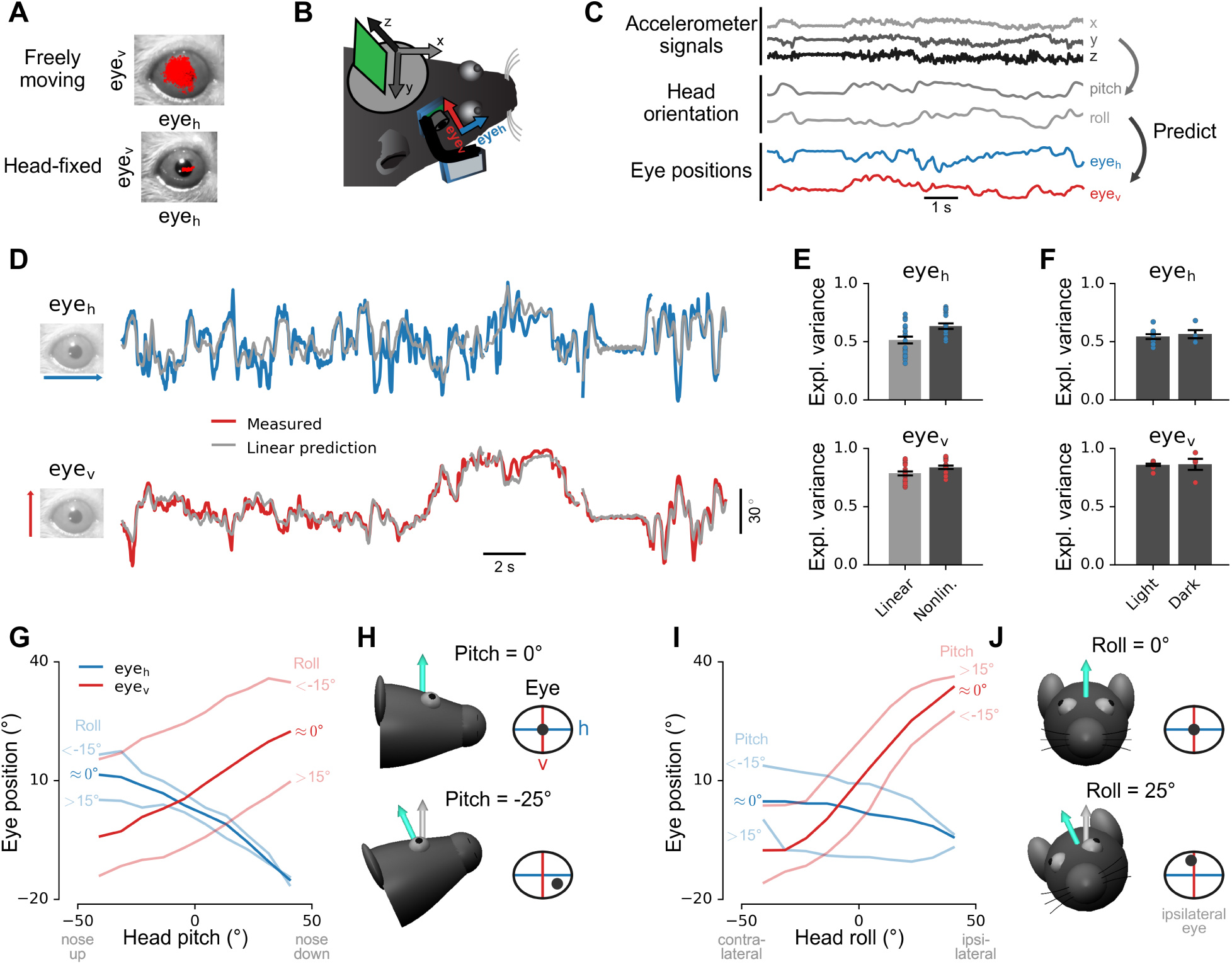
Systematic relationships between eye position and head orientation in freely moving mice. (A) Measured eye positions (red dots) in a freely moving mouse (top) and in the same mouse during head fixation on a cylindrical treadmill (bottom). (B) Method for simultaneous recording of eye position and head acceleration. (C) Head orientation (pitch/roll) was computed from low-pass-filtered head acceleration signals and was used to train models to predict eye position (arrows). (D) Measured eye positions compared to head-orientation-based predictions of a linear model. Model parameters were determined using training data different from the test data shown here. (E) Fraction of variance in eye position explained by head orientation, based on cross-validated predictions of linear (light gray) or nonlinear (dark gray) model. Top, horizontal eye position. Bottom, vertical eye position. 20 recordings in 3 mice (*n* = 8, 6, 6 in mouse 1,2,3 respectively, 10 minutes each). (F) Fraction of variance in eye position explained by head orientation using the nonlinear model in light (*n* = 10 recordings) and dark (*n* = 4 recordings) conditions (all sessions from one mouse, 10 minutes each). (G) Horizontal (blue lines) and vertical eye position (red lines) as a function of head pitch. Dark and pale lines show interaction with head roll: " 0°", 15° < head roll *<* 15°; "<-15°", head roll *<* 15°; ">15°", head roll *>* 15°. (H) Illustration of systematic dependence of horizontal and vertical eye position on head pitch, for pitch = 0°(top) and pitch = 25°(bottom). Eye and eye coordinate system (h/v) rotates with head. (I,J) The same as in G,H but as a function of head roll, and with dark and pale lines showing interaction with head pitch.

First, we examined the accuracy with which head pitch and roll predicted eye position (Figure 6C). Regression models based on these two variables were able to capture a large fraction of the variation in horizontal and vertical eye positions (Figure 6D,E; see also Methods) For a simple linear model, cross-validated explained variance between measured and predicted eye position was 52% for horizontal and 79% for vertical eye position; for a nonlinear model (see Methods), explained variance was 64% and 84% respectively (Figure 6E). Results were consistent over multiple months within and across mice, as indicated by the stability of regression model weights (Figure S6A). Explained variances were comparable in light or dark conditions (Figure 6F; see Methods for details), indicating that this effect of head orientation on eye position was driven by vestibular input or efferent signals rather than visual input (Andreescu et al., 2005; Oommen and Stahl, 2008).

Model predictions of eye position based on head pitch and roll were significantly more accurate for vertical than horizontal eye position (Figure 6E; Wilcoxon signed-rank test, *P*=2 *•* 10^*−*6^ for linear model, *P*=1 *•* 10^*−*5^ for nonlinear model). We wondered if the horizontal eye position might be more affected than the vertical by correlated movements across the two eyes independent of head orientation and used a dual-camera system to monitor both eyes simultaneously (Figure S6 and Movie S1). We then trained predictive models on data from each eye individually and found that the interocular error correlation (the correlation between variability in eye position not explained by pitch and roll of both eyes) was significantly stronger for horizontal than vertical eye position (*cc* = 0.72 horizontal, *cc* = 0.11 vertical; Wilcoxon signed rank text, *P* = 0.002; *n* = 10 recordings in one mouse, 10 minutes each).

We further asked if rotational head movements around the gravity axis (yaw), which are not well captured by the head-mounted linear accelerometer, might also account for the apparently weaker dependence of horizontal than vertical eye position on head orientation. To test this, we added a gyroscope to the implant (see Methods). Including rotations about the yaw axis increased the variance explained by the linear and nonlinear models by approximately 0.10 in horizontal and 0.02 in vertical eye position (Figure S6D), confirming some contribution of head yaw movements to prediction of horizontal eye position. The linear weights associated with the yaw signal were also remarkably similar across recordings (Figure S6B). In three recordings in the mouse with dual-camera implants and gyroscope, we found that interocular error correlation in the horizontal direction increased from 0.72 (head pitch/roll only) to 0.78 (including yaw as covariate) with no change in interocular error correlation in the vertical direction (0.12). Thus, coupled variation of eye position unexplained by orientation or rotation occurs primarily in the horizontal direction and may be caused by correlated eye movements not dependent on head movement, for example during resetting eye movements (van Alphen et al., 2001; Stahl, 2004) or continuous drift towards a resting eye position (van Alphen et al., 2001; Wallace et al., 2013).

Figure 6G,I summarizes the effects of head orientation on eye position. Both horizontal and vertical eye position varied systematically (and approximately linearly) with head pitch (Figure 6G) while vertical eye position was primarily affected by head roll (Figure 6I), consistent with reports in head-fixed mice (Oommen and Stahl, 2008) and freely moving rats (Wallace et al., 2013). Predictions of horizontal eye position were further improved by incorporating head yaw signals from a head-mounted gyroscope (Figure S6E). These results indicate that eye position is closely linked to head orientation in freely moving mice, even in the dark and even when the animals are exploring objects in enriched environments (Movie S6).

### Rapid eye movements are strongly linked to head movements in freely moving mice

We next investigated the relationship between eye and head dynamics. Angular head velocity was measured with the head-mounted gyroscope described above. Eye speed measurements taken around the time of increases in head rotation speed revealed a close correspondence between the temporal profiles of eye movements and head movements (Figure 7A), with eye movements on average in opposite directions to head movements (Figure 7B). These results are consistent with the observed dependence of eye position on head orientation (Figure 6G-J) and with the expected effects of the vestibulo-ocular reflex (VOR; Stahl, 2004).

**Figure 7:**
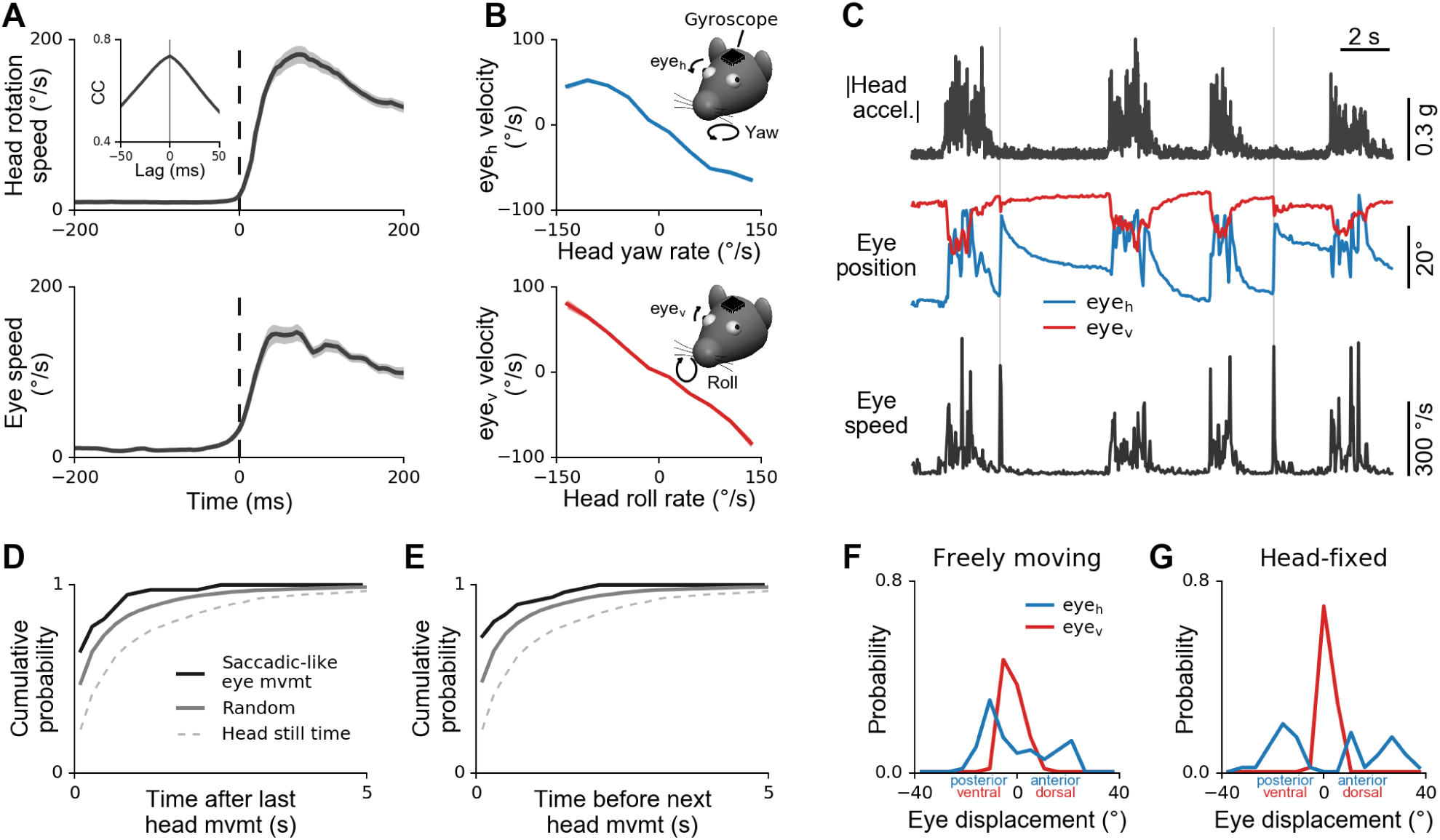
Coupling between eye movements and head movements. (A) Dynamics of head movement (top) and eye movement (bottom) during head movement initiation. Head rotation speed was measured using a gyroscope attached to the implant; eye speed computed from pupil positions. Traces were aligned to the onset of head movement (rotational speed ≥ 15°/s with at least 0.5 s of no movement before onset). Plots show mean ± SEM for *n* = 160 head movement events in one mouse, recorded in 14 different 10-minute sessions across more than 4 months. Inset shows average cross-correlation between head and eye speed; note peak at zero time lag. (B) Top: average horizontal eye velocity as a function of head velocity about the yaw axis. Directions as shown in inset. Bottom: average vertical eye velocity as a function of head velocity about the roll axis. In both directions, eye movements counteract head rotations. Plots show mean ± SEM (smaller than line width). Same dataset as in A. (C) Rapid eye movements occurring in the absence of head movements. Example traces showing magnitude of head acceleration computed from accelerometer signals (top), horizontal/vertical eye positions (middle), and eye speed computed from eye positions (bottom). Saccadic-like eye movements occurring in the absence of head movements (thin vertical lines) were identified by detecting eye movements with peak eye speed *>*250°/s which occurred when head movements were below a fixed threshold (0.0625 g). (D) Cumulative probability of the time between detected saccadic-like eye movements and the preceding head movement (solid dark line). For comparison, cumulative probability is also shown for simulated data (solid gray line) with the same saccadic-like eye movement rate but with saccades occurring at random times within the recorded head-still times (dashed line). Saccadic-like eye movements were significantly more likely to occur soon after a head movement than would be expected by chance (Kolmogorov-Smirnov test, *P* = 3.5 · 10^*−*8^). Same dataset as in Figure 6 (20 recordings in 3 mice, 10 minutes each). (E) Same as in D but for the time between saccadic-like eye movements and subsequent head movements. Saccadic-like eye movements were significantly more likely to occur just before a head movement than would be expected by chance (Kolmogorov-Smirnov test, *P* = 2.2 · 10^*−*7^). (F) Changes in horizontal and vertical eye position from 20 ms before to 20 ms after the time of peak speed in saccadic-like eye movements. Saccadic-like eye movements tend to be larger horizontally than vertically. Same dataset as in D and E. (G) Same as in F but for mice head-fixed on a cylindrical treadmill (4 recordings in 2 mice, 10 minutes each).

Despite this close overall coupling between head and eye movements, saccadic-like (*>*250°/s, see for example Sakatani and Isa, 2007) eye movements were occasionally observed in the absence of head movements (Figure 7C), occurring at an average rate of 0.044 Hz during head-still times. Moreover, these saccadic-like eye movements were not uniformly distributed during head-still times, but were significantly more likely to occur right before or after a head movement (Figure 7D,E). Saccadic-like eye movements were qualitatively similar in freely moving animals and head-fixed mice. Figure 7F,G shows the distribution of eye displacements in the horizontal and vertical direction for saccadic-like eye movements in freely moving and head-fixed mice, respectively. Interestingly, the largest eye displacements in freely moving mice were observed in the horizontal direction, consistent with the pattern in head-fixed animals. In freely moving mice, however, the range of horizontal eye displacements was slightly reduced (median movement magnitude 9.9°and 17.7°, respectively; Wilcoxon test, *P* < 3 *•* 10^*−*8^), perhaps reflecting greater reliance on head movements for gaze shifts in freely moving animals.

We conclude that eye movements are generally closely coupled to head movements in freely moving mice. Occasionally, the eye moves in the absence of head movement - but this typically happens just before or after a head movement. Together with the previous observation that average eye position is closely linked to head orientation even during active exploration, these results indicate strong interactions between eye and head movements at both fast and slow timescales in freely moving mice.

### Visual cortex activity is modulated by head movements in the dark

When combined with an implanted neural recording device, the head-mounted camera and motion sensor make it possible to investigate how brain activity is modulated during natural movements in freely moving mice. Previous work has indicated that locomotion modulates visual cortical activity in head-restrained mice (Niell and Stryker, 2010; Saleem et al., 2013; Fu et al., 2014; Reimer et al., 2014). We wondered whether head movements would evoke distinct patterns of activity in visual cortex (V1), given that V1 receives substantial vestibular input accompanying eye movements (Rancz et al., 2015; Vclez-Fort et al., 2018) along with inputs from many other non-primary sensory areas (Leinweber et al., 2017). We measured pupil, whisker pad, and head movements along with neural activity in single cells in V1 while animals freely explored a circular environment (Figure 8A) in the dark (to exclude the possibility of uncontrolled visual inputs during head movement). We tracked the body of the mouse and excluded periods of gross body movement (*≥* 1 cm/s) to analyze head movements that were not accompanied by locomotion (Figure 8B and Methods).

**Figure 8:**
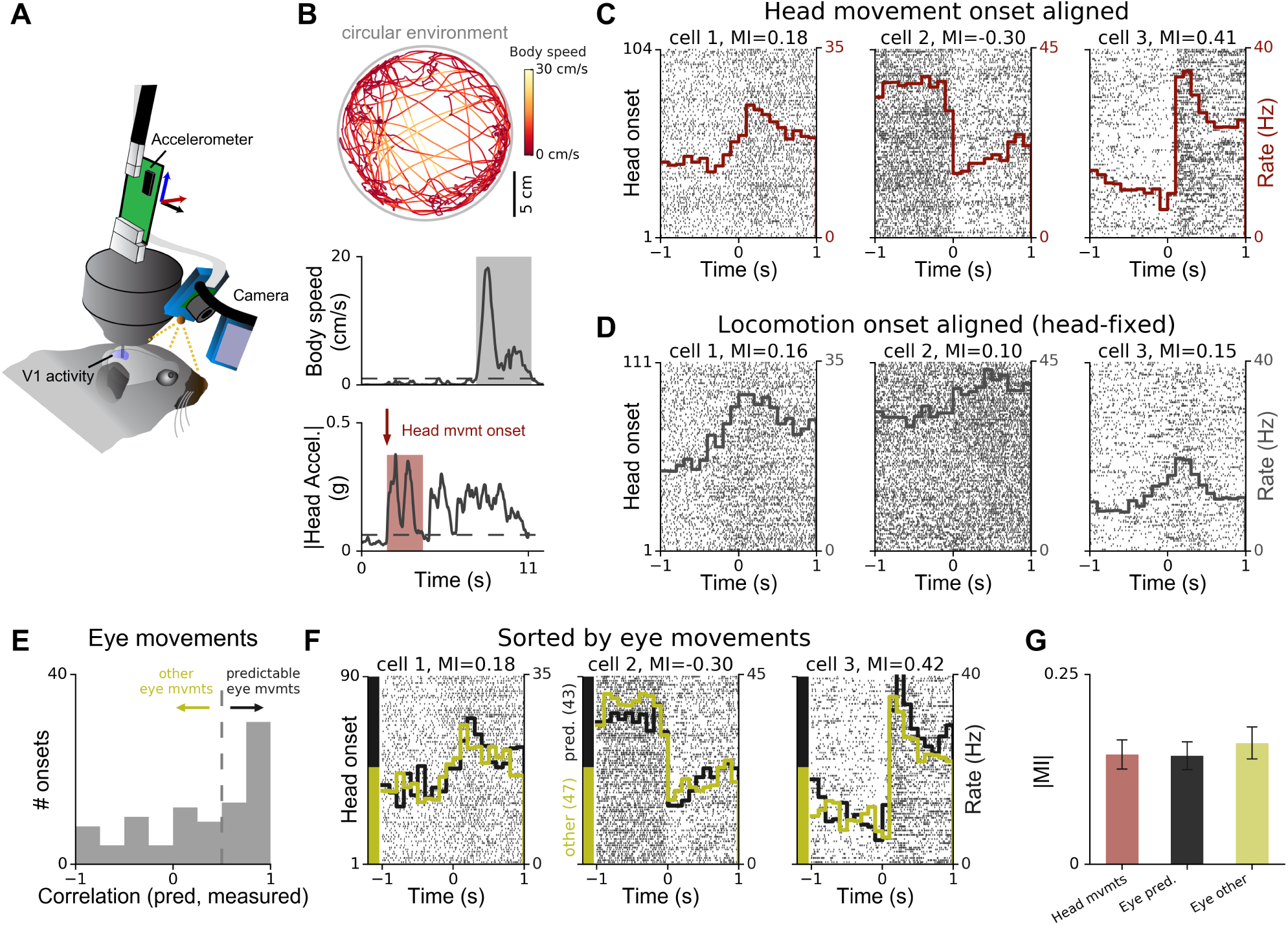
Head movement-related modulation of firing in visual cortex. (A) Chronic tetrode implant, head-mounted camera system, and head-mounted accelerometer were used to record neural activity in primary visual cortex (V1), eye positions, whisker pad movements, and head movements while mice explored a circular environment in the dark. (B) Top, body position and speed were tracked using an external camera. Middle, periods when body speed exceeded 1 cm/s (gray rectangle) were excluded from consideration in order to focus on head movements occuring without locomotion. Bottom, a head movement episode (red area) was defined as a period when body speed was less than 1 cm/s and head movement was above threshold (dashed line) following at least 0.5 seconds below threshold (before head movement onset). (C) Raster plots for three simultaneously recorded V1 cells, showing spike times relative to head movement onset. Rasters are displayed vertically according to onset index (i.e., time order) within recording (left axis). Red histograms show the average spike rate across trials (right axis). For all three cells, firing rate was significantly modulated by head movement (Wilcoxon signed-rank test, pre versus post movement onset; **P* <* 0.001). (D) Raster plots for the same cells as in C but aligned to locomotion onset (threshold 1 cm/s) for mouse head-fixed on a cylindrical treadmill. (E) Division of eye movement onsets into those well-predicted by a model based on head orientation (correlation ≥ 0.5) and other eye movements (correlation ≤ 0.5). (F) Raster plots and firing rate histograms for the same three cells as in C, for the two types of eye movement onsets shown in E. Spike train data same as in C, but including only head-movement onset events for which the eye movement could be reliably extracted. Rasters are grouped vertically by eye movement onset type as indicated by colored y-axis bars ("predictable", black; "other", yellow). Spike rate histograms shown overlaid using same color convention. (G) Summary of modulation indices (MI; see text) for V1 activity when aligned to head movement onsets, eye movement onsets that were predictable from head acceleration, or eye movement onsets that were not predictable from head orientation. Plot shows mean *±* SEM across 16 recordings (20-40 minutes each) in 3 mice (74 cells with at least 2 spikes per second).

Activity was tightly locked to head movement onsets in many visual cortical cells. In total, 55% (41/74) of V1 cells were significantly modulated by head movement (see Methods for details on spike sorting and data extraction). We observed both increases and decreases in firing rate even for simultaneously recorded cells (Figure 8C). To quantify the movement-related response modulation of individual cells, we computed a modulation index MI=(Post-Pre)/(Post+Pre), where Post and Pre are the mean firing rates for 1 s after and before movement onset, respectively. As shown for the three simultaneously recorded cells in Figure 8C and D, V1 response modulation at the onset of head movements without locomotion in unrestrained mice could be similar to or different from V1 response modulation at the onset of locomotion in the same animals head-fixed on a cylindrical treadmill. There was no significant correlation between the firing patterns of 74 V1 cells recorded in both conditions in 3 different mice (Wald test, *P* = 0.18; Figure S7A). This observation suggests that head movements can affect firing rates of visual cortex neurons independently of locomotion.

Head movements were tightly coupled to eye movements in freely moving mice (Figure 7A). To disentangle the effects of these variables on V1 responses, we first asked if eye movements differentially affected V1 activity in the dark. We extracted the first eye movement in the period around head movement onset by measuring optical flow of the pupil edges in the dark (Methods and Figure S8). We then used the eye position models described above to predict eye movements from head accelerometer data. Head/eye movement onsets were classified by whether the initial eye movement was in the same direction as the predicted eye movement based on head movement (correlation *≥* 0.5) or in another direction (correlation *<* 0.5). Approximately half of eye movements were in the direction predicted by the model based on head movement (Figure 8E). Sorting the trials of the cells in Figure 8C according to whether or not the eye movement was predictable from the head movement did *not* reveal any systematic differences between the two conditions (Figure 8F). There was no significant difference in absolute modulation indices between unsorted and sorted conditions (Figure 8G) across all recorded cells with at least 20 trials in all conditions (N=37; Wilcoxon signed-rank test, *P* = 0.8). Indeed, MI values for spike trains aligned to head movement onsets accompanied by either "predictable" or "other" eye movements were both statistically indistinguishable from MI values obtained when the same spike trains were aligned to onsets of the head movements regardless of eye movements (Figure S7B,C; *P* = 0. 06 for "head mvmt" vs "predictable", *P* = 0.15 for "head mvmt" vs "other"; Wilcoxon signed-rank test). These results suggest that eye movements did not have a differential impact on the observed modulation of V1 responses, beyond that predicted by head movements alone.

We next asked whether whisker movements differentially affected modulation of neural responses in V1. Whisking was not as strongly coupled to head movements as eye movements. We aligned V1 recordings to the onset of whisking and observed that the magnitude of modulation was generally reduced (Wilcoxon signed rank test, *P* = 0.001) compared to alignment to head movement. Furthermore, alignment to the onset of whisking in the absence of head motion resulted in an even greater reduction (Wilcoxon signed rank test, **P* <* 3 *•* 10^*−*6^; Figure S7D-F). This indicates that head movements modulate V1 activity more strongly than whisking movements in most recorded cells.

We conclude that head movements modulate V1 activity in freely moving mice, even in the dark and in the absence of locomotion. Moreover, while head, eye, and whisker movements are coupled in freely moving mice, modulation of V1 activity by head movements cannot be fully explained in terms of coupled eye movements or whisking alone.

## Discussion

The mouse is a prominent animal model in neuroscience, but behavioral monitoring in freely moving mice has been limited by the absence of video tracking methods in head-centered coordinates. To overcome this limitation we developed a miniature ultra-lightweight head-mounted video camera system, and combined it with movement sensors to monitor multiple behavioral variables including pupil size and eye position, head, whisker pad and body movements, and integrated it with a chronic multielectrode implant to record neural activity in freely moving animals (Figure 1). The camera system is stable, enabling precise and continuous monitoring of behavioral variables and minimizing the amount of postprocessing required to extract the variables of interest. Inter-frame image movement was less than 1 pixel (corresponding to about 40 *µ*m) in about 99% of all video images, even when the mice were grooming, exploring complex environments, or interacting with objects in the environment (Figure 3 and Movie S2). Crucially, mouse behavior was similar with and without the camera system, allowing accurate monitoring of pupil size, eye position, whisking, and other variables during natural behaviors. The operation of the camera system did not affect the quality of simultaneous electrophysiological recordings.

This new head-mounted camera system significantly expands the range of scientific questions that can be addressed in freely moving mice. Ethological studies could reveal the precise characteristics of behavior such as eye movements, whisking, and other motor outputs. Sensory neuroscientists could use the system to validate experimental results obtained under conditions of head or body restraint - while directly studying sensory processing under more natural conditions. Studies of non-sensory brain areas, including associative and motor areas, could identify sources of behavioral variability that drive neural activity but have been previously hard to measure. Mouse models of disease could be examined to establish or to exclude deficits in eye movements, whisking or other motor outputs.

Here we have shown that the head-mounted camera system can provide new insights into the relationships between eye, head, and whisking movements and neural activity in freely moving mice. In many animals, eye and head movements are intimately related and both are used for orienting gaze towards salient objects (Land, 2015). However, very little is known about their coordination in mice, even though this information could provide important general insights into how non-foveate animals use vision during natural behavior. We observed prominent changes in the distributions of both head orientation and eye position in different behavioral states in freely moving mice. When we quantified this relationship using predictive models, we discovered that a large fraction of the variation in eye position could be predicted from head orientation, consistent with findings from a previous study in the freely moving rat (Wallace et al., 2013). Our results suggest that freely moving mice stabilize their gaze relative to the horizontal plane. Crucially, our data show that this gaze stabilization does not only happen on average but also at a fine temporal resolution (Figure 6 and Movie S6), and therefore may play an important role in mouse vision. We also found that the systematic relationships between eye position and head orientation were preserved across months, across mice, and in the dark as well as the light, suggesting that head-orientation-related changes in eye position are driven by vestibular rather than visual input (Oommen and Stahl, 2008).

While models based on head orientation and rotational head movements were able to explain most variation in eye position particularly in the vertical direction, there was still considerable unexplained variance in the horizontal direction (about 10-50%). By using two head-mounted cameras we found that horizontal eye positions not explained by head orientation were strongly correlated across both eyes, even after taking into account rotational movements of the head. Whether these correlations resulted from resetting eye movements not locked to head movements (e.g. van Alphen et al., 2001) or active shifts in gaze will need to be determined in future work. Most of the present experiments were done in a circular environment without salient visual objects. However, in enriched environments it appeared that mice did not orient their eyes towards objects even when they actively explored them (Movie S1). Moreover, even saccadic-like eye movements occurring without a coincident head movement were significantly more likely to occur just before or just after a head movement than would have been expected by chance. Future experiments might use the camera system to investigate whether freely moving mice encountering highly salient or moving visual objects produce more eye movements that are not coupled to head orientation or head movements. Furthermore, monitoring not only the eyes but also the environment using a head-mounted camera facing outward without IR mirror (Movie S1) will help to clarify the link between head and eye movements and visual inputs.

We also demonstrated how the camera system can be combined with motion sensors and chronic neural recording devices to discover new relationships between motor-related variables and neural activity in the visual cortex. About 55% of V1 cells were modulated by head movements in the absence of locomotion, even in the dark, i.e., in the absence of any visual input. Both enhancement and suppression of firing were seen, even for cells recorded at the same time. These results were not explained by variations in eye movements or whisking. Recent work has demonstrated that locomotion can modulate activity in sensory cortex (Niell and Stryker, 2010; Schneider et al., 2014). For example, in mouse primary visual cortex, neural responses are generally enhanced when head-fixed animals run on a treadmill compared to when they are stationary (Niell and Stryker, 2010); in contrast, in primary auditory cortex, neural responses are typically suppressed by locomotion (Schneider et al., 2014). We measured changes in neural activity in primary visual cortex either during head movements in the absence of locomotion when the mouse was freely moving, or during locomotion when the mouse was head-fixed on a cylindrical treadmill. We found that the directions of modulation in the same V1 neuron could be different for locomotion-related and head-movement-related responses, and that there was no significant correlation between the two types of movements. These results demonstrate that modulation of early sensory cortical areas by motor outputs is both more general (i.e., occurring for many forms of movement) and more specific (i.e., manifested differently for different forms of movement) than previously thought. Future work will be needed to identify whether the movement-related signals are used for suppression of sensory coding during self-generated movement (e.g. saccadic suppression, Duffy and Burchfiel, 1975), for the computation of the mismatch between sensory input and expected input (Keller et al., 2012), or for the integration of sensory inputs with signals related to spatial navigation (Saleem et al., 2013). We anticipate that important progress can be made by combining our method with new tools for virtual reality in freely moving animals (Stowers et al., 2017; Del Grosso et al., 2017) to provide both detailed behavioral and stimulus control.

The new system is open-source and we provide all required software and design files. To our knowledge this is the first open-source head-mounted video tracking system for small laboratory animals. The system uses widely available components (e.g., camera sensor, single-board computer and connectors) or 3D-printable parts (camera holder), and the total cost is low (see parts list) which should further promote its adoption. Moreover, this ultra-lightweight system could be easily adapted for use in larger animals, such as rats, ferrets, and monkeys. At the moment the system is tethered, but especially in larger animals it is possible to add batteries to power the system so it can be used in conjunction with wireless recording methods (Fan et al., 2011; Yin et al., 2014). In the mouse, a major challenge remains the weight of the combination of headposts, cameras, parts for neural recordings, batteries and wireless transmitters, but technical developments in miniaturizing all these components might make entirely wireless head-mounted neural recording and behavioral monitoring systems feasible in the near future. Furthermore, the system is modular and could be integrated with alternative methods for recording neural activity, such as high-density silicon probes (Jun et al., 2017) or head-mounted fluorescence microscopes (Zong et al., 2017), and/or combined with technologies for optogenetic manipulation of neural activity during behavioral monitoring (Wu et al., 2015).

Because the position of the camera and mirror can easily be customized, the view can be modified to include other variables of interest. For example, a small modification to the arm holding the mirror is suffi cient to provide a detailed image of the pinna (Movie S1) to provide insights into how pinna movement contributes to the processing of incoming sounds, e.g., during sound localization, in freely moving animals. The camera could also be used to monitor the movement of single whiskers in head-centered coordinates, as opposed to the whisker pad movements tracked in the current study, without the need for external tracking cameras, computation of absolute position in space, or attachment of markers to single whiskers (Voigts et al., 2008; Roy et al., 2011; Nashaat et al., 2017). Finally, the camera system can also be used to capture images of the nose, mouth, and/or paws, to monitor how mice interact with their environment when they explore novel objects (see Movie S1 for a mouse interacting with Lego and foraging) and during social behaviors such as mating and fighting. Thus, the system has the potential to greatly increase the range and scope of experimental questions that can be addressed about natural behaviors in freely moving mice and other small laboratory animals.

## Acknowledgements

We thank Jakob Voigts for providing the drive parts for the initial experiments, and John Stahl and Takashi Kodama for their advice on tracking eye position in the dark. The authors are also grateful to Stephen M. Town and Nicholas A. Lesica for their comments on the manuscript. This work was supported by the Biotechnology and Biological Sciences Research Council (BB/P007201; J.F.L.); Action on Hearing Loss (G77; J.F.L.); the Gatsby Charitable Foundation (M.S.) (GAT3212, J. O’K); the Simons Foundation (SCGB323228; M.S.); the UCL Excellence fellowship (J.P.); and the Wellcome Trust (090843/C/09/Z, 090843/D/09/Z, J. O’K). J.O’K. is a Wellcome Trust Principal Research Fellow (203020/Z/16/Z).

## Author contributions

A.F.M and J.P. conceived the project and designed and performed the experiments; A.F.M. designed and constructed the experimental setup and the camera system; A.F.M. performed computational and statistical analyses and prepared figures; J.F.L. supervised animal procedures; M.S. and J.F.L. supervised analysis; All authors discussed the results; A.F.M. and J.P. drafted the manuscript; All authors contributed to the writing of the manuscript.

## Methods

### Contact for Reagent and Resource Sharing

Further information and requests for resources and reagents should be directed to and will be fulfilled by the Lead Contact, Jennifer F Linden (j.linden©ucl.ac.uk).

## Experimental Model and Subject Details

### Animals

Experiments were performed on male C57Bl/6J mice (Charles River) for visual cortex recordings and male C57Bl/6J and CBA/Ca mice (Charles River) for the sound experiment. After surgical implantation of chronic implants for neural recordings, mice were individually housed on a 12-h reversed light-dark cycle (lights off at 12.00 noon). Water and food were available ad libitum. All experimental procedures were carried out in accordance with a UK Home Offi ce Project Licence approved under the United Kingdom Animals (Scientific Procedures) Act of 19J86.

## Methods Details

### Surgical procedures

For chronic implants, we used custom tetrode hyperdrives with 8-16 individually movable tetrodes, constructed according to a published design (Voigts et al., 2013). Tetrodes were made from HM-L coated 90% platinum/10% iridium 17 *µ*m diameter wire (California Fine Wire). A miniature male connector (Omnetics NPD-18-DD-GS) was attached to the front of the drive body (see "Construction of the lightweight camera system") for connection of the camera system during behavioral experiments.

Mice aged 58-65 days were anaesthetized with 1-2% isoflurane and injected with analgesia (Carprofen, 5 mg/kg IP). Ophthalmic ointment (Alcon) was applied to the eyes, and sterile saline (0.1 ml) injected SC as needed to maintain hydration. A circular piece of dorsal scalp was removed and the underlying skull was cleaned and dried. A custom machined aluminum head-plate was then cemented onto the skull using dental adhesive (Superbond C&B). A small craniotomy was made over the left primary visual cortex (V1) (2.5 mm lateral, 1 mm anterior to the transverse sinus). The tetrode drive was positioned above the craniotomy and fixed to the skull with dental adhesive. A pinhole craniotomy above the right prefrontal cortex contralateral to the tetrode implant for the ground screw (000-120 × 1/16, Antrin Miniature Specialties). The ground screw and implant were then secured with more dental adhesive and dental cement (Kemdent Simplex Rapid). Mice were allowed to recover from surgery for at least five days before experiments began.

### Neural recordings in head-fixed and freely moving mice

All experiments were conducted in a custom double-walled sound-shielded anechoic chamber. Animals became accustomed to handling and gentle restraint over two to three days, before they were head-fixed and placed on a custom styrofoam cylinder (20 cm diameter, on a ball-bearing mounted axis). After animals were head-fixed the headstage was connected to the implant and the camera holder was connected to the miniature connector on the outside of the implant, together with two cables from the headstage which provided power to the IR light-emitting diode (IR LED).

We confirmed that each tetrode recording site was in monocular V1 by presenting stimuli on a screen contralateral to the implant side and identifying the approximate receptive field position of recorded cells as described previously (Poort et al., 2015). Luminance of visual stimuli was calibrated using a luminance meter (Konica Minolta, LS-100). Running speed on the cylinder was detected with a rotary encoder (Kübler, 1024 steps per rotation) and single steps were extracted using a microcontroller (Arduino Uno), sent to the recording system as transistor-transistor logic (TTL) pulses and recorded along with neural data.

For experiments in freely moving mice, the implant was gently held while allowing the mouse to walk or run on a running wheel, and headstage and camera system were connected as for the head-restrained experiments. The animal was then released into a circular environment for experiments in the freely moving condition. Two different circular environments were used. The first environment (diameter 30 centimeters) consisted of white plastic material. Eight LED lights (ULT300, Digital Daffodil) combined with custom cut light diffuser sheets (Perspex) were used to provide homogeneous lighting which facilitated tracking of the eye (see "Extraction of pupil positions from camera images"). For the sound experiment (Figure S5) a loudspeaker was mounted 1 meter above the center of the environment (see "Sound presentation"). The second environment (diameter 22 centimeters) consisted of black plastic materal with a semi-transparent perspex floor to allow reliable tracking of body position using an external camera from below (see "Analysis of head movement onsets"). This second environment was used to perform recordings in the dark (Figure 8).

Neural activity was recorded with a 32-channel Intan RHD 2132 amplifier board (Intan Technologies) connected to an open-ephys acquisition board (Open Ephys) via an ultralightweight flexible serial peripheral interface cable (Intan Technologies). Data were sampled at 30 kHz and saved to disk for off-line analysis.

### Electrophysiological data analysis

Electrophysiogical recordings were analysed off-line using Bayesian spike-sorting techniques (Sahani, 1999). To detect action potentials the common median reference was subtracted across channels (Rolston et al., 2009) with subsequent high-pass filtering with a cutoff of 600 Hz, and action potentials were detected by finding time points exceeding 3.5 times the standard deviation of the noise. Action potentials were automatically clustered. Single units or small clusters of neurons were accepted only if the spike-sorter reported both false-negative and false-positive rates below 5 %. Clustered units were verified manually and units were classified as single-unit (SU) if fewer than 0.5 % of the spikes occurred within the typical refractory period of a cortical neuron (*≤* 2 ms). All other units were deemed multi-units (MUs).

The effect of the head-mounted camera system on neural recording quality was assessed using raw broadband signals and spike units (158 SUs and 11 MUs). The power spectral density (PSD) of broadband signals was estimated using Welch’s method with a 2 s long Hann window and 1 s overlap. For each condition, the PSD of all electrode channels was computed separately and the log-scaled PSDs averaged afterwards to yield a single estimate of the PSD (Figure 2C). To quantify the difference across all recordings, we computed the PSD ratio between segments with camera on and off (10 minutes each) recorded during the same session without disconnecting the neural recording headstage (Figure 2D-F). The order of the two conditions was balanced across sessions to reduce potential effects of behavioral changes during each session (e.g., mice typically explored the environment more during the early part of the recording). Within-condition variability for the implant-only condition was estimated by computing the standard deviation of PSD ratios for different non-overlapping 60 s segments from the same recording. The signal-to-noise ratio (SNR) between spikes and high-pass filtered electrode signals (Figure 2E) was computed as the power of the electrode channel of each tetrode with maximum depolarization, and the noise power extracted from electrode signals between spikes (with a 2 ms margin around spikes). All data recorded during the same session were spike sorted together to avoid the need to manually register spike clusters between conditions.

To compute the power in the local field potential (LFP), raw traces were first bandpass filtered at 2 - 10 Hz (low-frequency LFP in Figure 5) or 10 - 20 Hz (higher-frequency LFP in Figure 5) using a zero-phase fourth-order Butterworth filter with subsequent squaring of the filter output. The resulting estimate of the LFP power was smoothed with a normalized Gaussian window with a standard deviation of 2 seconds before computing the correlation with pupil dilation (Figure 5D). For visualization, LFP power was normalized such that low-frequency LFP power had a mean value of 1 (Figure 5A,C).

### Construction of the lightweight camera system

We used a commercially available camera module (Adafruit 1937) with an Omnivision OV5647 sensor capable of 640 × 480 pixels per frame at up to 90 Hz. The CMOS camera sensor has dimensions of 8.2 mm x 11.3 mm x 4.8 mm and weighs 0.5 grams (including suspended part of the cable). The infrared (IR) filter was removed to allow monitoring of behavioral variables in dark conditions using IR light. The sensor was attached to the neural implant using a custom camera holder. The camera holder consisted of a 3D printed frame with clips for holding the camera sensor (Figure S1). A lightweight 21G steel cannula (length *∼* 2 cm) for holding the IR mirror (Qioptiq, NIR-Blocking Filter, Calflex-X) was bent by about 75°in the middle, inserted with one end into a hole in the frame and fixed with epoxy resin (Araldite Steel). The mirror was cut to size 7 mm x 7 mm and attached to the cannula via a 3D printed holder. This enabled fine adjustment of the mirror relative to the camera sensor by moving the mirror along the cannula, rotating the mirror around the cannula, and also by further bending the cannula. A miniature connector (Omnetics NSD-18-DD-GS) for mounting the camera system to the implant was attached to the back of the 3D printed holder base using super glue (Loctite Power Flex Gel). After final adjustment of the mirror, either during surgery or during head-fixation of the animal on a running wheel (see "Neural recordings in head-fixed and freely moving mice"), the cannula and the mirror holder were permanently fixed using a thin layer of strong epoxy resin (Araldite Rapid). STL and OpenSCAD source files for the camera and mirror holders will be made freely available.

Illumination of the camera’s field of view, including eye and whisker pad, was provided by a small IR LED (Vishay VSMB2943GX01) mounted to either the bottom or the side of the camera holder, depending on the angle between camera sensor, mirror, and implant. The IR LED was powered by the headstage via two 36AWG wires and a small-package current-limiting resistor (Farnell Multicomp, 100 - 180 Ohm, metric package size 3216). Custom cut gold pins (RS Pro Male and Female Solder D-sub Connector Contact, 481-493 and 481-500) soldered to the wires and the headstage allowed quick and stable connection during experiments. All parts, including weight and estimated cost, are summarized in Table S1. An example camera holder is shown in Figure S1B.

### Interfacing with the camera

The camera was connected to a single-board computer (Raspberry Pi 3 model B, Raspberry Pi Foundation) with ARM architecture and VideoCore 4 graphics processing unit (GPU). Data from the camera were read out with custom software using the Multi-media Abstraction Layer (MMAL) API (Broadcom Europe). Because miniature cameras such as the one used for the head-mounted system do not typically provide additional output signals to synchronize frame acquisition, we used the following approach to avoid dropped frames during recording and to obtain time stamps that were precisely synchronized with neural recordings. First, each frame was annotated with a time stamp from the GPU immediately after acquisition. Once the frame was received and decoded by the custom software, a TTL signal pulse was sent to the recording system using the general-purpose input/output capabilities of the single-board computer. The difference between the acquisition and TTL signal time stamps was saved to a separate file for post-hoc alignment of TTL time stamps and neural data. Communication between the computer for recording neural data and the single-board computer for controlling the camera was done via ethernet using the ZeroMQ messaging library (http://zeromq.org/). Automatic starting/stopping of the camera system was controlled using a custom plugin for the open-ephys recording system (http://www.open-ephys.org/). Code for frame acquisition, TTL time stamp generation and alignment, and the plugin for controlling the camera will be made freely available.

Figure S2 demonstrates precision of aligned time stamps for a blinking LED (Vishay TSAL4400, typical rise/fall time 800 ns) recorded under the same conditions as the behavioral data in the experiments, for different video resolutions and frame rates. The LED was driven by a microcontroller (Teensy 3.2, PJRC) and the same signal was sent to the recording system. The pixel corresponding to the maximum LED intensity was identified and LED onset times were extracted from the pixel intensity trace by thresholding at 0.5 full intensity.

### Detection of camera image movements

For each recording, movement of the camera image was detected by selecting a region of interest (ROI) that contained a part of the neural implant (inset in Figure 3A). A correlation-based algorithm (Dubbs et al., 2016) was used to detect movements between the average ROI (averaged across all recorded images) and the ROI for each video image. Using the average ROI as reference image ensured that whisker or hair movements on single images did not have an impact on the overall detection performance. Images with changes in brightness exceeding three standard deviations were excluded from the analysis to remove periods when the camera view was blocked, e.g., during grooming. On average only 0.6% and 0.2% of the camera images were removed from the freely moving and head-restrained recordings based on this criterion, respectively.

### Extraction of pupil positions from camera images

In order to perform tracking of pupil positions, it was necessary to remove bright regions from the camera image resulting from reflections of the illumination IR LED on the cornea. Therefore, contiguous bright regions on the recorded camera frames were detected by thresholding, and a binary mask was generated. Thresholds were manually selected for each session to include the major IR LED reflections. The original frame and the binary mask were used to estimate the values of masked pixels using non-texture image inpainting (März, 2011). An ellipse was fitted to the processed frame by thresholding, contour extraction, and least-squares ellipse fitting (Fitzgibbon et al., 1999). Contour extraction thresholds were manually adjusted for each session and only ellipses with mean pixel intensities below a user-defined threshold and with areas above another user-defined threshold were kept to reduce false positive rates. Thresholds were selected based on a small number of eye frames (*≤* 2%) randomly selected from the whole recording. Finally, ellipses were manually verified using custom software including a graphical user interface. Ellipse fitting code will be made freely available.

In behavioral experiments where we tracked the eye position in the dark, we administered an eye drop of physostigmine salicylate (0.1-0.2%) 30 minutes in advance to limit pupil dilation (see for example Oommen and Stahl, 2008).

### Extraction of whisking pad movement from camera images

Movement of the whisker pad was extracted by selecting a rectangular region of the camera image containing the whisker pad. Dense optical flow was computed (Farnebäck, 2003) and the average optical flow across all pixels was used as a measure of whisker pad movement in horizontal (related to azimuth) and vertical (related to elevation) directions. All analyses in this study were based on horizontal movements.

We compared whisker pad movements recorded using the head-mounted camera (60 Hz) to data recorded simultaneously using an external camera (100 Hz) from above while the mouse was head-fixed. The head-mounted camera was able to capture important aspects of whisking including the whisking frequency and fluctuations in whisking envelope (Figure S3).

In some experiments described here (e.g., Figure 5, Figure S5), the camera system was operated with a frame rate of 30 Hz, and therefore whisker pad movements were measured only up to 15 Hz. In principle, however, the camera could be run at 90 Hz frame rates to capture more detailed aspects of whisking (e.g., whisker angles), using more sophisticated algorithms to extract these parameters at high frame rates (Perkon et al., 2011).

### Extraction of head orientations from accelerometer signals

Gravity components in the accelerometer signals were estimated by low-pass filtering each channel with a zero-phase second-order Butterworth filter with a cut-off frequency of 2 Hz (Pasquet et al., 2016). Pitch, defined as the angle between the naso-occipital axis and the horizontal gravity plane, was extracted by computing the angle between the gravity vector and the y/z plane with normal vector **e**_*x*_= (1,0, 0)^T^. Roll, defined as the angle between the interaural axis and the horizontal gravity plane, was extracted by computing the angle between the gravity vector and the x/z plane with normal vector **e**_*y*_= (0, 1, 0)^T^.

To compute head orientation maps (Figure 4F,I and Figure 5F) the low-pass filtered accelerometer signals were transformed into spherical coordinates (with elevation angle Θ and azimuthal angle Φ). A 2D histogram of head orientation vectors with a bin size of 5°for both elevation and azimuth was computed on the unfolded sphere. In order to visualize the histogram on a sphere, the number of samples within a each bin was normalized by the corresponding quadrangle area. Normalized histogram data were color-coded on a logarithmic scale.

### Behavioral segmentation

Behavioral states were segmented using a semi-automatic classification algorithm. In a first step, about 1 - 2 hours of video recorded using external CMOS cameras (The Imaging Source, 20-50 Hz frame rate) were annotated manually for each mouse and for each condition ("Implant+cam" and "Implant" in Figure 4). Only behavioral segments with a duration of at least 2 seconds were assigned a behavioral state.

The behaviors that categorized were "grooming" (G), "eating" (E), "quiescence" (Q), and "active exploration" (A). Grooming comprised different stereotypical movements, e.g., movement of the forepaws over the nose and muzzle, strokes of forepaws across vibrissae and eye, and strokes with the hindleg. These movements were typically periodic and therefore easily distinguishable from the other behaviors. Eating was identified during chewing on seeds added to the environment. As chewing was also evident as artifacts on electrode channels we used this information during manual annotation but not during automatic segmentation. Because the sessions in which seeds were added to the environment were not balanced across conditions, we accounted for this during the analysis shown in Figure 4E,H by assigning the mean value across sessions with seeds to those without seeds. Periods when the mouse was still for at least 2 seconds were classified as quiescence and periods when the mouse was exploring the environment and not grooming or eating were classified as active exploration.

We found that segmentation based on the time-domain accelerometer signals (Venkatraman et al., 2010; Dhawale et al., 2017) resulted in relatively low accuracy of identification of the behaviors described above. We therefore developed an algorithm performing segmentation in the frequency domain that considerably increased accuracy compared to segmentation based on time-domain signals (Figure S4C,D). The algorithm worked as follows: accelerometer signals (Figure 4B) were transformed into a spectro-temporal representation using a short-time Fourier transform (STFT) with a Hann window of length 2 s and a window shift of 40 ms. At each time step, the log-scaled magnitude of the transformed accelerometer signals was recast as a single vector containing data from all accelerometer channels. The middle point of the window was used as reference point for the annotated behavioral category. A Multilayer Perceptron (MLP) with one hidden layer (N=100 hidden units with rectified-linear activation functions) was then fit to the data. The network was trained using the backpropagation algorithm and the weights were optimized using a stochastic gradient-based solver with adaptive momentum estimation (Kingma and Ba, 2014) via the sklearn Python package (Pedregosa et al., 2011).

We evaluated the prediction performance of the model using cross-validation. That is, the data set was divided into 4 parts, model parameters were estimated leaving out one of the parts, and the predictive quality of the model fit was evaluated on the part left out. This procedure was repeated leaving out each of the 4 parts in turn and the prediction accuracy averaged to yield an estimate of the goodness-of-fit of the model. The confusion matrices in Figure 4C and supplemental Figure S4 show the cross-validated true positive rate computed from the manually annotated data ("Human observer") and the prediction of the model.

To assess the differences between occupancies of the different states in the two experimental conditions ("Implant+cam" and "Implant" in Figure 4E,H) we computed the least absolute deviation (L1 norm) between the distributions for both conditions. To confirm the significance of this difference, we used a permutation test. A null distribution was generated by shuffl ing "Implant+cam" and "Implant" condition labels across recording sessions. This approach ensured that any significant differences from the null distribution could be attributed to the presence of the camera rather than time of the recording session (see Figure S4B,C). The permutation procedure was repeated 10000 times, and a *P*-value was generated by computing the fraction of permutations with least absolute deviations larger than the value computed on the original data set. The same permutation procedure was used to determine the significance of the difference between body speed distributions in the active state (Figure 4G,J). Mean and variance of head orientations (Figure 4F,I) were computed using a permutation test for the difference in circular mean and variance as test statistic, respectively.

## Sound presentation

Broadband noise burst stimuli (50 ms, 50 or 55 dB SPL, noise bust rate 0.5 Hz or 1 Hz) were generated using custom software, converted to an analog signal (HDSPe AIO, RME), amplified (RB-850, Rotel), and delivered via a loudspeaker (XT25TG30-04, Tymphany) mounted about 1 meter above the circular environment. Sound pressure levels of the acoustic stimuli were measured (40BF 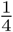 in free-field microphone and 26AC preamplifier, GRAS) and calibrated to the center of the circular environment. In the experiments shown in Figure S5, recordings with and without acoustical stimulation were interleaved (up to 5 minutes each, total duration 30 minutes) during periods when the animal was quiescent and immobile.

## Prediction of eye position using head orientation

Pupil positions were extracted from video data (sampled at 42-60 Hz) as described in "Extraction of pupil positions from camera images". Only time points at which the pupil could be detected were included in the analysis and no smoothing was applied for the analysis. For visualization, extracted eye position and pupil dilation traces were smoothed using a 3-point Gaussian window with coeffi cients (0.072, 0.855, 0.072). Head pitch and roll were computed from signals recorded using the 3-axis accelerometer (sampled at 7500 Hz) integrated into the neural recording as described in "Extraction of head orientations from accelerometer signals"

For each pupil position *p_i_, i* = 1, 2*, …, N*, the most recent history of each signal within a time window of 500 ms was recast as vector **u**_*i*_, **v**_*i*_for pitch and roll, respectively. Linear interpolation was used to find the pitch/roll at time lags −500, −475, −450, …, 0 ms.

Two different models were trained using the resulting data. The linear model assumes that pupil positions are related to the pitch and roll via

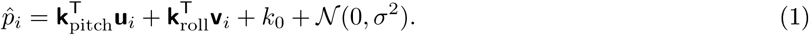

The linear weighting vectors **k**_pitch_ and **k**_roll_, and the offset term *k*_0_ were found using a Bayesian method for determining the relevance of inputs, known as Automatic Relevance Determination (ARD) (MacKay, 1996). Because the relation between accelerometer signals and pupil position can potentially be nonlinear we also tested a Multi-Layer Perceptron as described in Results.

In some experiments we added a lightweight gyroscope sensor (MPU-9250, InvenSense, San Jose, US) to measure angular velocity, including rotations about the yaw axis. The sensor was calibrated using a stepper motor (Adafruit 324) and a contact tachometer (DT-2235B, Lutron Electronic). To approximate angular yaw position we convolved the velocity signal with an exponential decay function with time constant *τ* = 1 s and extended the models to also included the recent history of angular positions (Eq. 1 and Figure S6F).

The prediction performance of the different models was evaluated using cross-validation as described above (but with *n* = 5 fold). Similarity between predicted and measured eye positions was quantified using the coeffi cient of determination 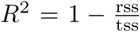 where rss is the residual sum of squares and tss is the total sum of squares.

In behavioral experiments where we tracked the eye position in the dark ("Extraction of pupil positions from camera images"), we typically recorded 2-3 segments (10 minutes each) before administration of an eye drop of physostigmine salicylate and one recording with eye drop (10 minutes) about 30 minutes after administration. About 20 minutes after the dark recording the pupil size was too small to allow for reliable tracking. This procedure was repeated on four different days in one mouse resulting in 10 recordings with light on and 4 recordings in the dark.

## Analysis of head movement onsets

Data for analysis of head movement onsets was collected while mice were exploring a circular environment (see "Neural recordings in head-fixed and freely moving mice"). The bottom of the circular environment consisted of an acrylic sheet that allowed reliable tracking of the mouse’s body using a camera placed below the environment, even in the presence of headstage and camera cables. Head movements were extracted from accelerometer signals by subtracting the gravity components (see "Extraction of head orientations from accelerometer signals"). The magnitude of head movements was computed as

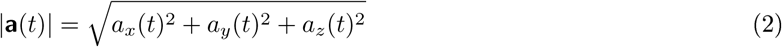

where *a*_*x*_, *a*_*y*_, and *a*_*z*_ are the head acceleration components along x, y, and z channels of the accelerometer, respectively, sampled at time step *t*. The magnitude was smoothed using a low-pass filter with a cut-off frequency of 2 Hz and thresholded using a fixed threshold across all mice and recordings (0.0625 g). Positive threshold crossings were classified as head movement onset if the smoothed magnitude of the accelerometer signals was (i) below the threshold for at least 0.5 s before and (ii) above the threshold for at least 0.5 s after the threshold crossing (Figure 8B). Moreover, movement onsets during locomotion periods (body speed *≥* 1 cm/s) were excluded from the analysis in Figure 8. Onsets of whisker pad movements and locomotion were computed in the same way as head movement onsets but whisking thresholds were selected separately for each mouse and the minimum duration above threshold was 0.1 s to account for faster movements of whiskers. Because mice were only occasional running during each recording session, presumably due to the relatively small size of the circular environment, we computed locomotion onsets for mice running on a cylindrical treadmill (threshold 1 cm/s) in the dark.

For the analysis, spike times were aligned to head movement onsets for each recorded V1 cell. To quantify the extent to which head-movement-related activity modulated the activity of each cell, we computed a modulation index (MI) defined as

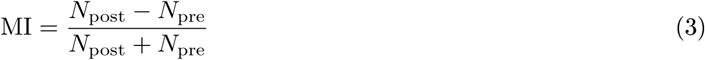

with *n*_pre_ and *n*_post_ denoting the average number of spikes 1 s before and 1 s after movement onset, respectively. MI values reported here were computed without subtraction of the baseline firing rate.

Because tracking of the pupil in the dark can be challenging due to increased pupil dilation (and because the effect of pharmacological intervention to reduce pupil dilation is not known, see "Extraction of pupil positions from camera images"), we extracted initial eye movements after movement onsets by measuring optical flow of the pupil edges in the dark. The region of the camera image containing the eye was filtered using a median filter with a window length of 15 pixels before computing optical flow of the pupil edges. This step ensured that movements of hair or IR LED reflections did not impair optical flow measurements. The flow for each pixel was computed using the same dense algorithm as for the whisker pad movements. To convert optical flow (measured in pixels per frame) to horizontal and vertical eye positions, we integrated the average flow for each dimension across time (i.e. frames). The integrated flow provides an approximation to initial eye movements after a head movement onset (but might diverge after some time due to potentially leaky integration of the flow measure). Comparing flow-based pupil positions to direct pupil fitting in dim light conditions (i.e. when the enlarged pupil was still possible to identify using ellipse-based pupil fitting), we found that analysis of optical flow of pupil edges yielded reliable estimates of eye positions after head movement onsets in the dark (Figure S8).

To test whether different types of eye movements have an effect on the observed head movement-related modulation of V1 firing, we divided movement onsets into two groups: head movement onsets that were consistent with predictions of eye positions based on the models described above, and onsets that were not consistent with model predictions. Because the observed modulations of V1 firing were fast (typically appearing less than 100 ms after the head movement onset) we used the x/y values of first peak of the measured and predicted eye movements as an approximation to the initial movement. This yielded one x/y pair for the measured and one x/y pair for the predicted trace following an onset. Only pairs with a maximum/minimum within 100 ms after the head movement onset were included in the analysis. The values in Figure 8E and Figure S8C show the correlations between both x/y pairs.

## Quantification and Statistical Analysis

Specifics on the statistical methodologies and software used for various analyses are described in the corresponding sections in Results, figure legends, Methods details, and supplemental figures. Statistical test results are described as significant in the text where *P* < 0.05.

## Data and code availability

Software to control the camera and to perform data extraction, along with 3D models for custom parts in the camera system, will be made available at http://www.gatsby.ucl.ac.uk/resources/mousecam/. Further data from this study are available from the corresponding authors upon reasonable request.

## Supplemental Figures

**Figure S1:**
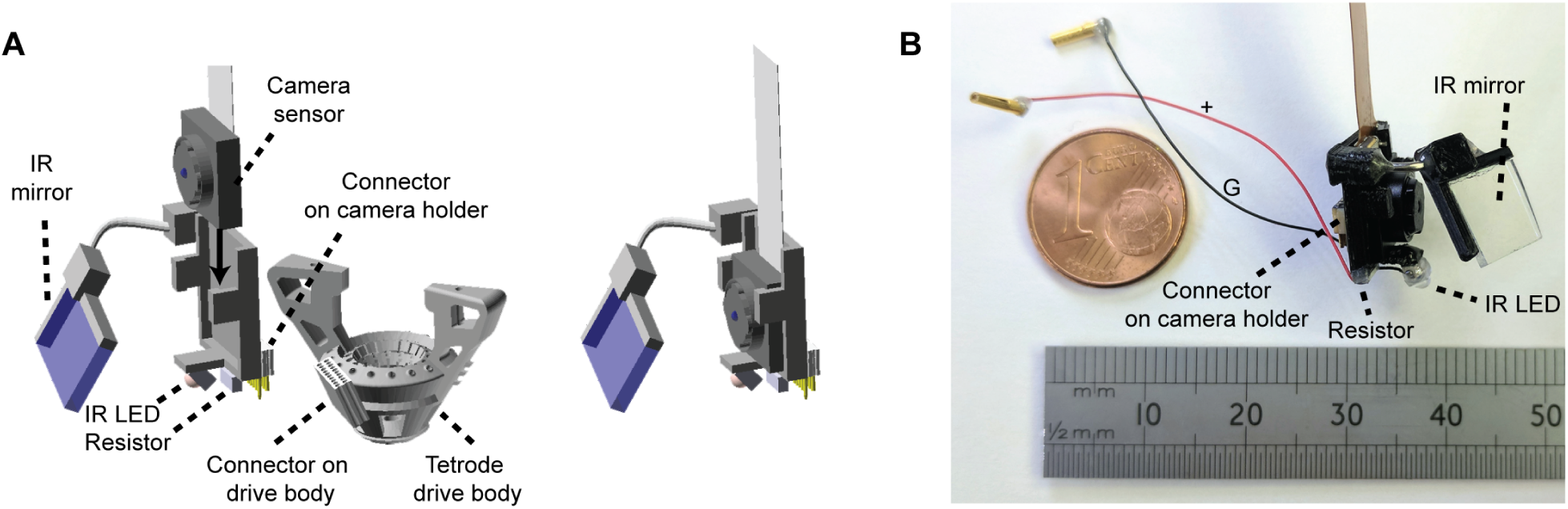
Design of the miniature head-mounted camera system. Related to Figure 1. (A) 3D computer-aided design (CAD) model showing the entire camera assembly, including camera sensor, camera holder, IR mirror, IR LED, miniature connector on the camera holder, and complementary connector permanently fixed to the drive body of the tetrode implant. The camera sensor is mounted on the camera holder via small clips on the sides of the 3D-printed holder. Connectors allow the camera system to be attached to the neural implant for recording sessions, and removed otherwise. (B) Example camera holder. Black and red wires are connected to ground (G) and positive (+) pins on the headstage (respectively), to provide power to the IR LED via the current-limiting resistor. Euro cent coin (left) and ruler (bottom) are shown for size comparison.

**Figure S2:**
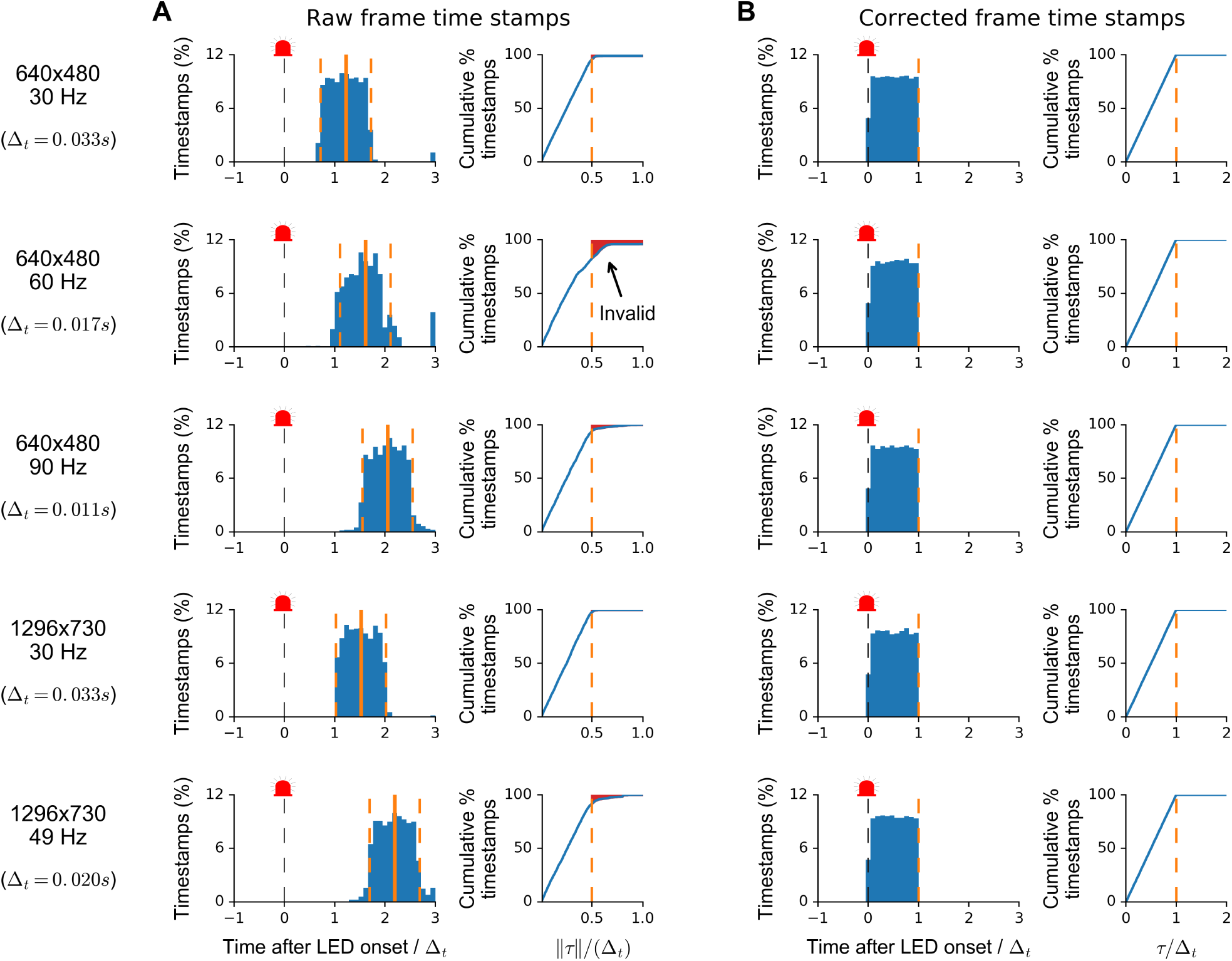
Correction of video timestamps. Related to Figure 1. (A) Distributions of raw frame timestamps (left) generated using the single-board computer controlling the head-mounted camera. Timestamps were normalized by the frame interval of the camera (∆_*t*_) and mean and 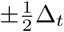 are indicated by the solid and dashed orange lines, respectively. The cumulative fraction of timestamps with time ±*τ* around the mean (right) reveals that not all timestamps were generated within the interval 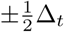 as indicated by the shaded red area. Therefore using the mean delay does not allow for reliable and precise synchronization of camera and neural data. Each row represents results for the video format shown on the left. (B) The same but after correcting video timestamps as described in Methods. All frame timestamps were within the frame interval [0, ∆_*t*_).

**Figure S3:**
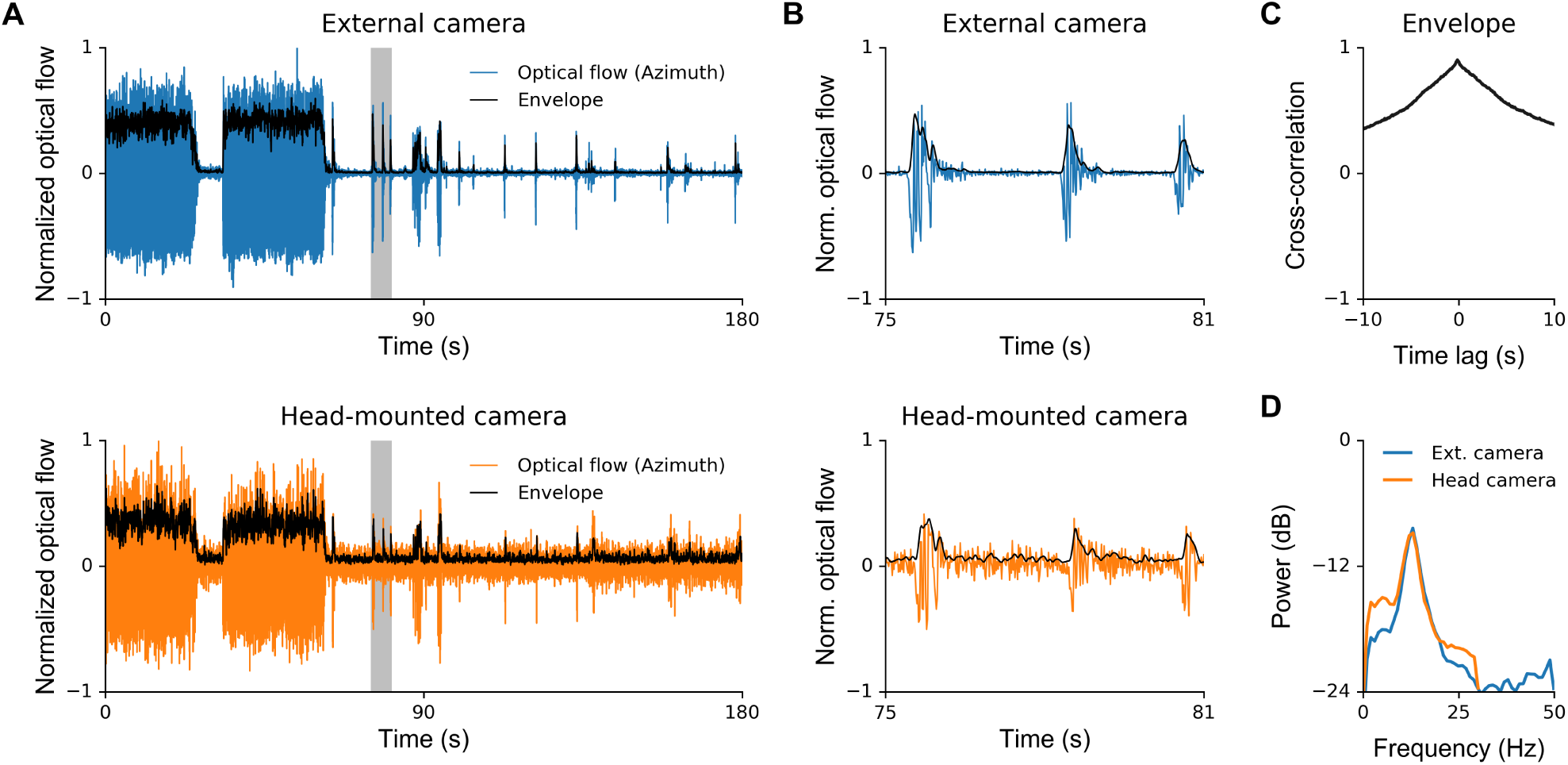
Correspondence between whisker pad movement estimates obtained from head-mounted and external camera images. Related to Figure 4. (A) Top: Optical flow extracted from external camera images of the whiskers viewed from above in a head-fixed mouse; sampling rate 100 Hz. Bottom: Optical flow extracted simultaneously from head-mounted camera images of the whisker pad; sampling rate 60 Hz. Optical flow was normalized to the interval [−1, 1]. Black lines indicate envelope obtained by low-pass filtering the magnitude of the optical flow with a causal filter with a cutoff frequency of 5 Hz. (B) Zoomed-in traces for gray shaded region in A. (C) Cross-correlation function between envelopes extracted from the external camera signal and the head-mounted camera signal in A. (D) Power spectral densities estimated from the optical flow signals in A.

**Figure S4:**
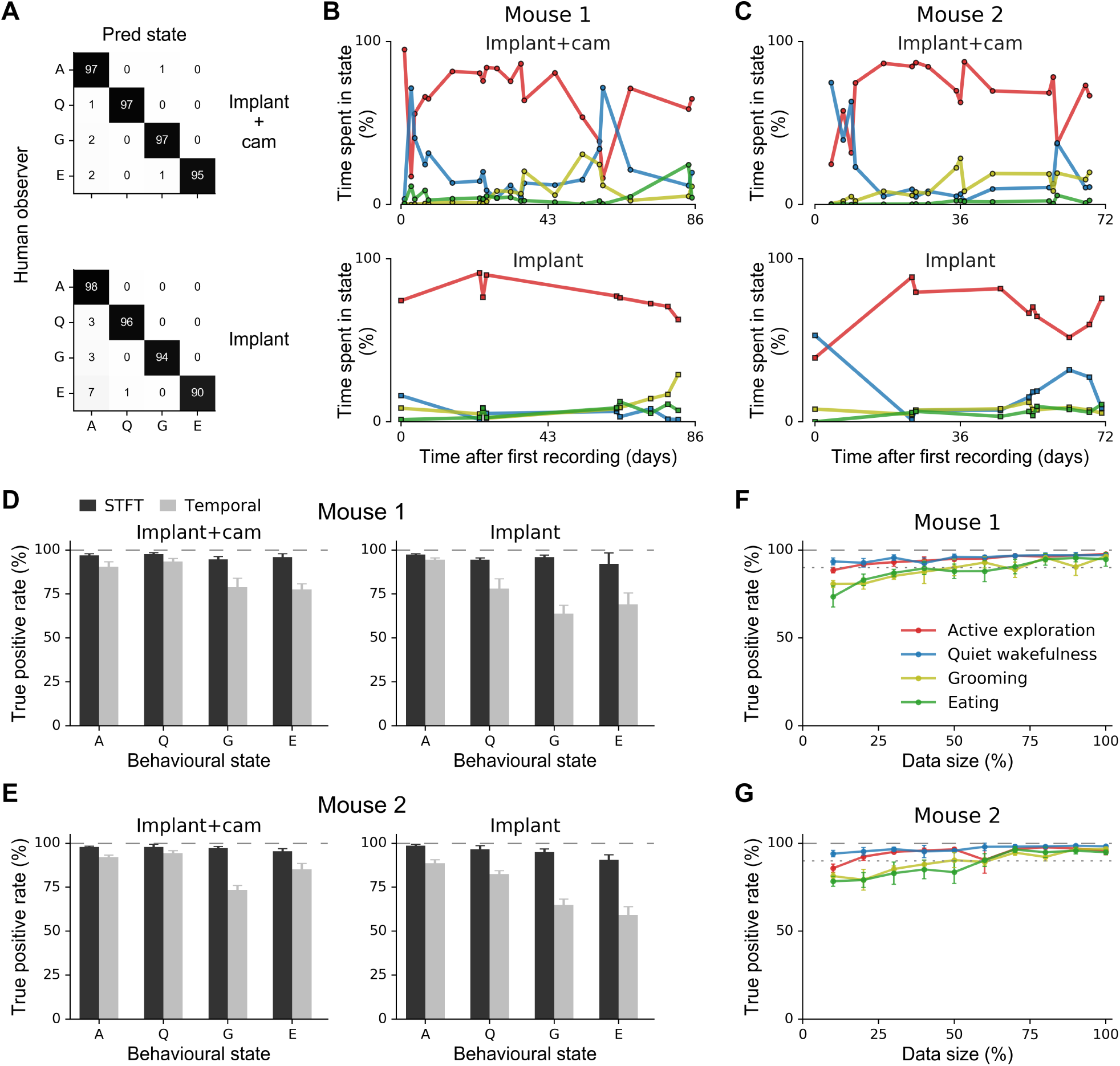
Details on behavioral segmentation. Related to Figure 4. (A) Confusion matrix showing cross-validated classification performance for second mouse. Top: Mouse with implant and camera. Bottom: Mouse with implant only. (B,C) Time spent in the different behavioral states as a function of days after first recording for the two mice. Top: Mouse with implant and camera. Bottom: Mouse with implant only. Same legend as in F. (D,E) Cross-validated segmentation accuracy (true positive rate) for the temporal accelerometer representation ("Temporal", gray bars) used in Venkatraman et al. (2010) and the spectral (short-term Fourier transform) representation proposed in this study ("STFT", black bars) for the two mice, both with and without camera. Total time of annotated data: "implant+camera", 90 minutes for mouse 1 and 60 minutes for mouse 2; "implant": 60 minutes for mouse 1 and 60 minutes for mouse 2. (F,G) Cross-validated segmentation accuracy as a function of training data size for the "implant + camera" condition. 100% corresponds to 58 minutes and 45 minutes of recorded data for mouse 1 and 2, respectively. Validation set sizes were 19 and 15 minutes for mouse 1 and 2, respectively, and validation was performed using the same 4-fold cross-validation scheme as in D and E. The results suggest that about about 75% of the annotated training data are suffi cient for behavioral segmentation with *>*90% accuracy (gray dotted line).

**Figure S5:**
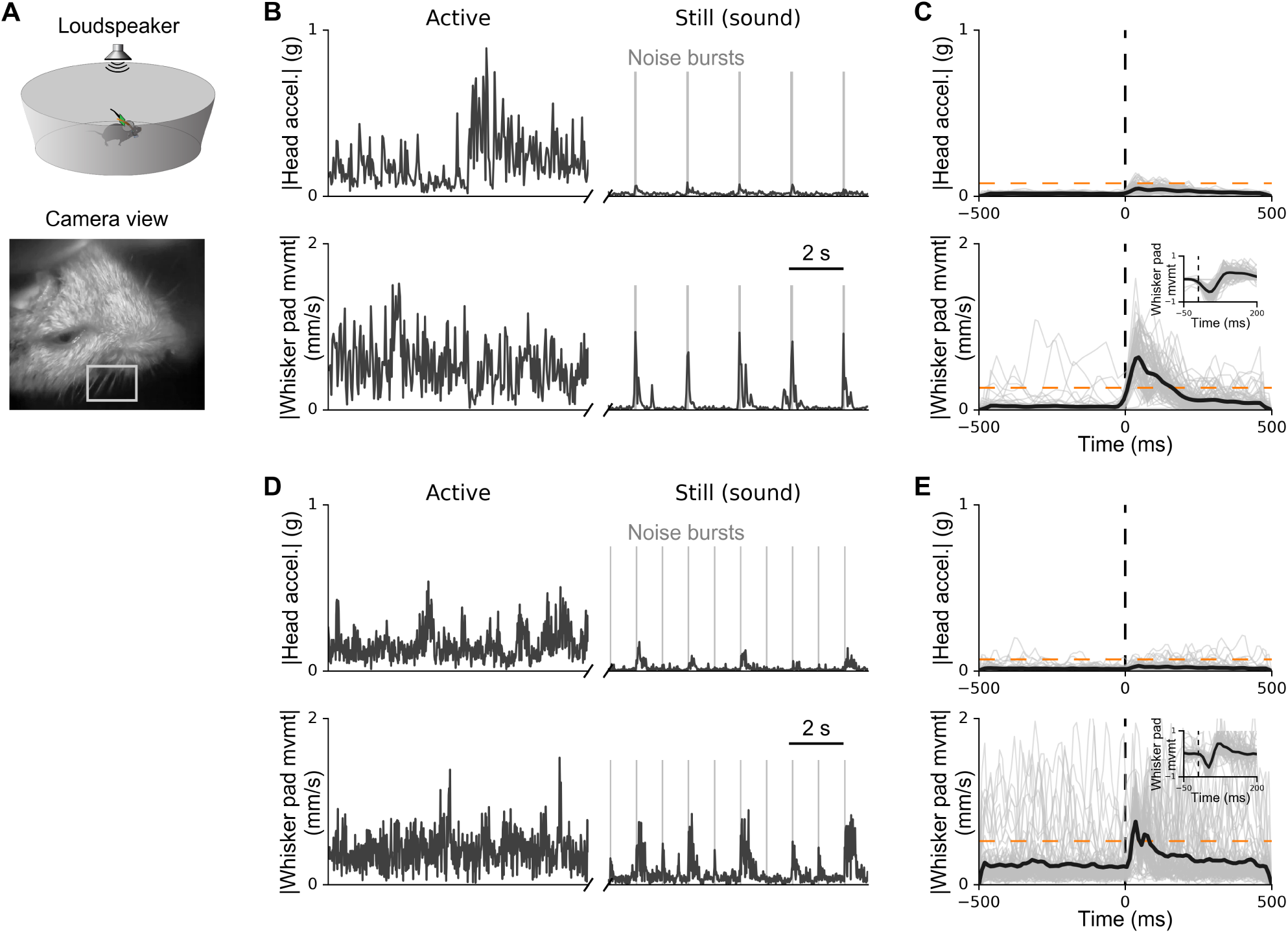
Sound-evoked whisker movements measured with the head-mounted camera. Related to Figure 5. (A) Top: Sounds were presented via a loudspeaker mounted 1 meter above the center of the circular environment. Bottom: Example frame from head-mounted camera focused on whiskers from above (gray rectangle). (B) Example traces showing head movement (top) and whisker pad movement magnitude (bottom) for an active period when the mouse was exploring the environment, and a still period when the mouse was immobile. During the still period, noise bursts (55 dB SPL, 50 ms) were presented every 2 seconds. Sound onset-triggered head movements (top) and whisker pad movements (bottom). The dashed orange line indicates one standard deviation for movements observed in the active period, for comparison. Sound-evoked whisker movements follow a stereotypical protraction/retraction movement pattern (inset). (D,E) The same as in B,C but for second mouse. Noise bursts (50 db SPL, 50 ms) were presented every second.

**Figure S6:**
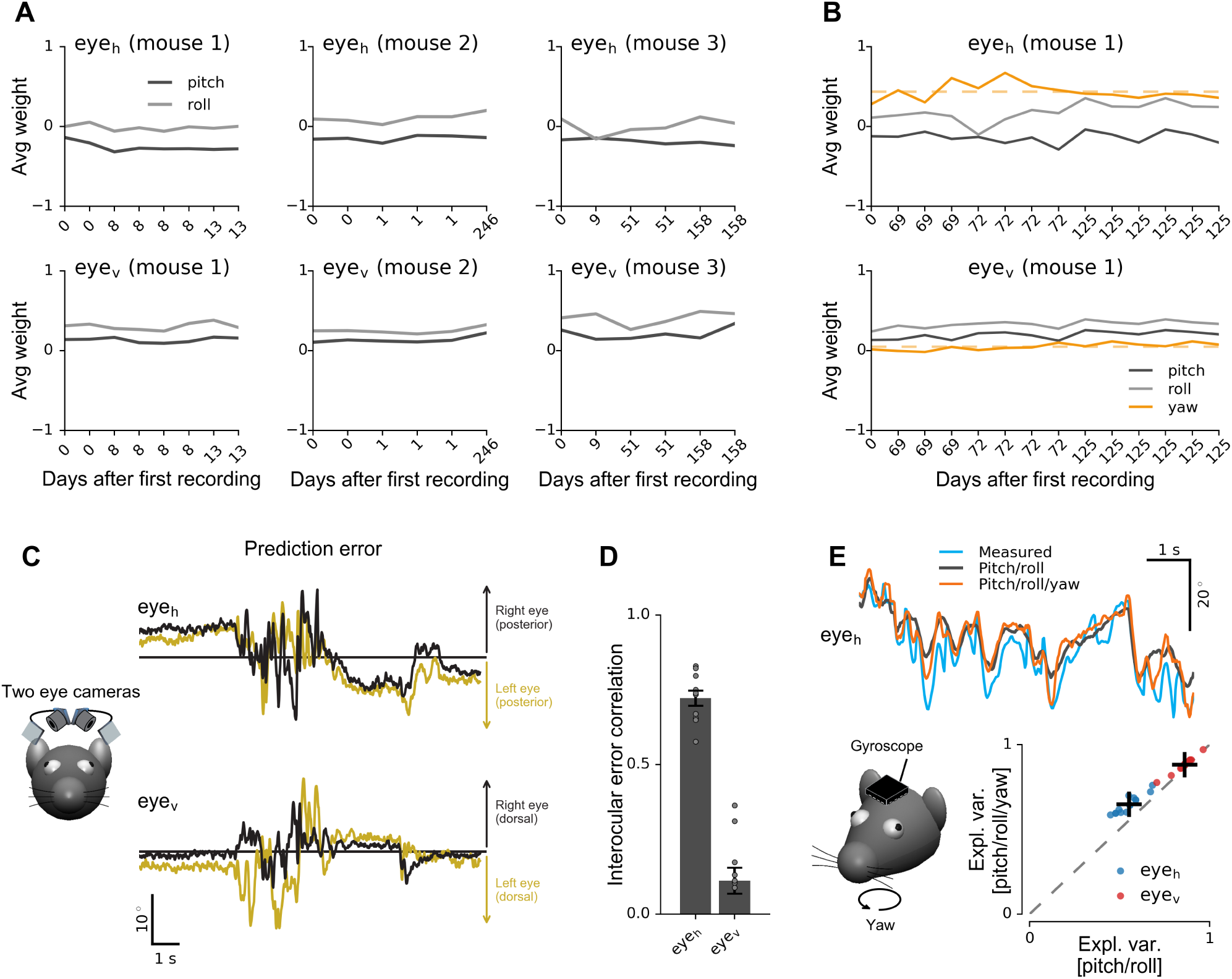
Details on prediction of eye position from head orientation. Related to Figure 6. (A) Average weights of linear model as a function of time after first recording, illustrating stability of estimated weights across recording sessions. Top and bottom rows show weights for horizontal and vertical eye position, respectively. Repeated recording days indicate multiple recordings (10 minutes each) on the same day. Same data as for the three mice shown in Figure 6E. (B) The same as in A but for 14 recordings in one mouse with accelerometer and gyroscope sensors (yaw). (C) Simultaneous monitoring of both eyes using two head-mounted camera systems in a freely moving mouse. Example traces show prediction errors of the pitch/roll-based nonlinear model for horizontal (top) and vertical (bottom) eye position. Yellow lines, left eye. Black lines, right eye. (D) Correlation between model prediction errors for the two eyes, for 6 experiments in one mouse. Results indicate that failures to predict horizontal eye position based on head orientation (pitch/roll) were strongly correlated between the two eyes. (E) In addition to head acceleration, rotations about the yaw axis were measured using a head-mounted gyroscope (bottom left). Example traces show predictions of a nonlinear model trained with pitch/roll (black) and pitch/roll and yaw rate (orange). Cross-validated prediction performance increased mostly for horizontal eye position (bottom right). Increases in explained variance were statistically significant for both horizontal and vertical eye position, and for both linear and nonlinear models (Wilcoxon signed-rank test, **P* <* 1 *•* 10^*−*3^; *n* = 14 recordings; 10 minutes each).

**Figure S7:**
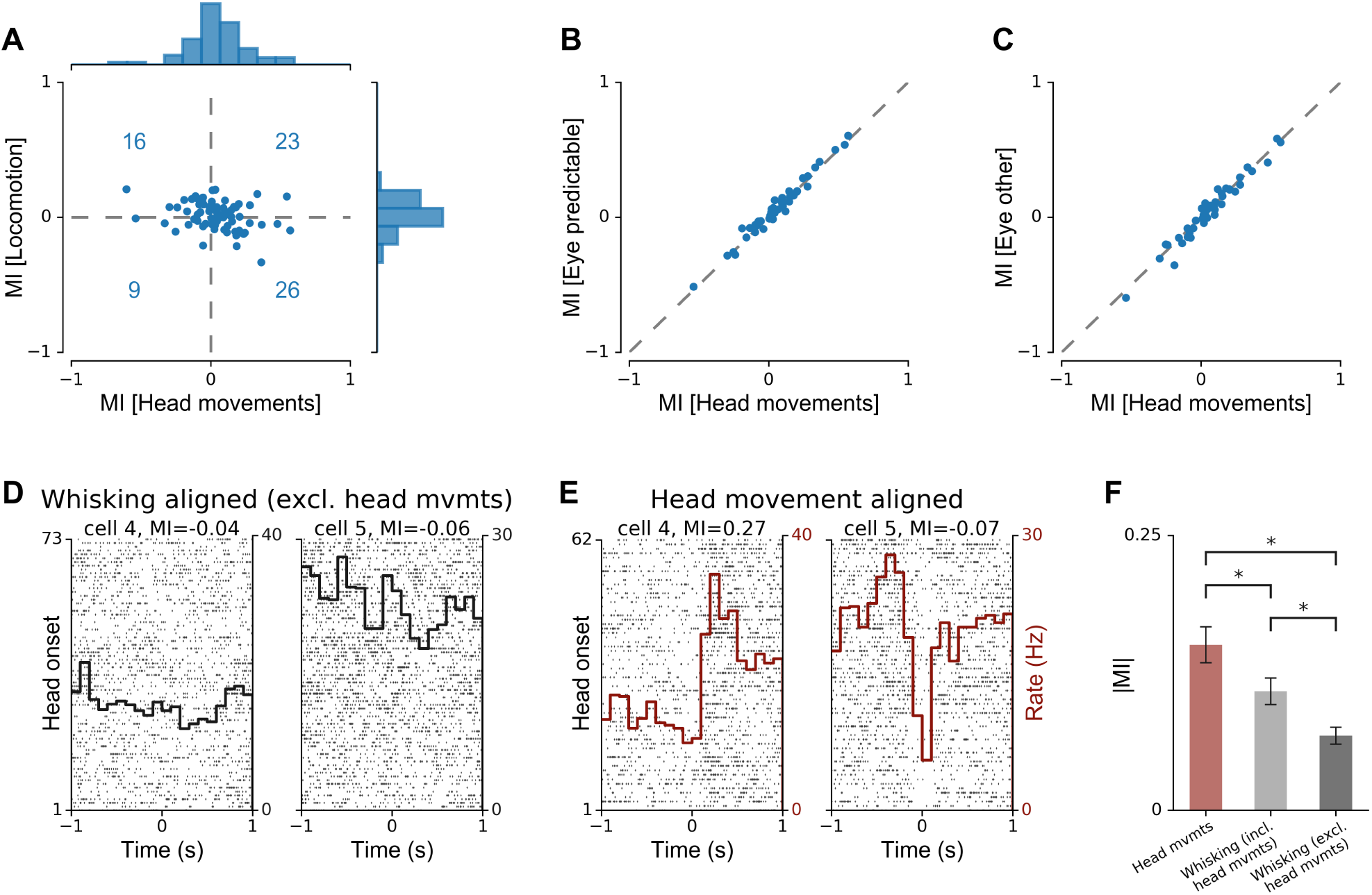
Modulation of V1 activity by head movements versus locomotion or whisking. Related to Figure 8. (A) Relationship between modulation indices (MIs) for V1 spike trains either recorded when mice were head-fixed on a cylindrical treadmill and aligned to locomotion onsets, or recorded when mice were unrestrained but immobile (body speed ≥ 1 cm/s) and aligned to head movement onsets. Plot shows data from all V1 cells recorded in 3 mice which had firing rates of at least 2 spikes/s in both conditions. Histograms show marginal MI distributions for head movement onsets (top) and locomotion (right). Blue numbers indicate absolute numbers of cells in each quadrant. (B) Comparison of the MI values for the same V1 spike trains for head and eye movements in which the eye movement was predictable from the head movement. (C) Same as B, but for head and eye movements in which the eye movement was not predictable from the head movement. (D) Spike rasters and rate histograms for recordings from two V1 cells, aligned to whisker pad movement onset after excluding periods of head movements. Other conventions as in Figure 8C. (E) The same cells as in H, but with spike times aligned to head movement onsets. (F) Comparison of absolute MIs for V1 activity when aligned to head movement onsets (red bar), whisking onsets when head movements are included (light gray bar), and whisking onsets excluding periods of head movement (dark gray bar). Plot shows mean ± SEM across 16 recordings (10-40 minutes each) in 3 mice.

**Figure S8:**
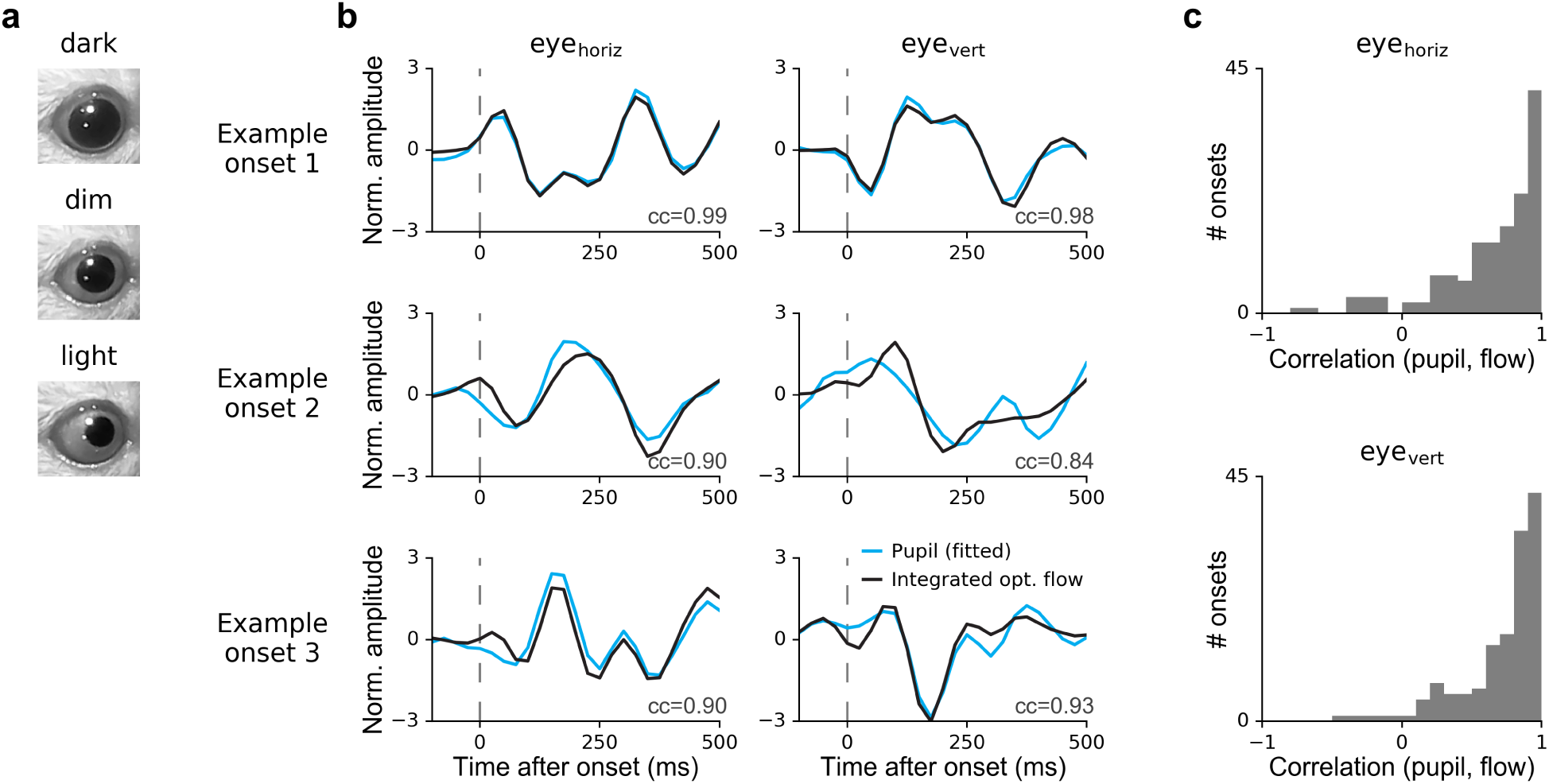
Optical flow-based pupil movement extraction. (A) Frames showing typical pupil dilation in dark (top), dim light (middle), and normal light (bottom) conditions. Dim and normal light conditions allowed direct fitting of pupil position. (B) Example horizontal and vertical eye movement traces extracted in the dim light condition, either by fitting an ellipse to the pupil (blue line) or by integrating optical flow (black line) of the pupil edge. Trends were removed by high-pass filtering. (C) Correlations between movement onset-triggered ellipse-fitted and optical flow-based pupil position traces for ten minutes of recorded data (144 onsets).

## Supplemental Tables

**Table S1:**
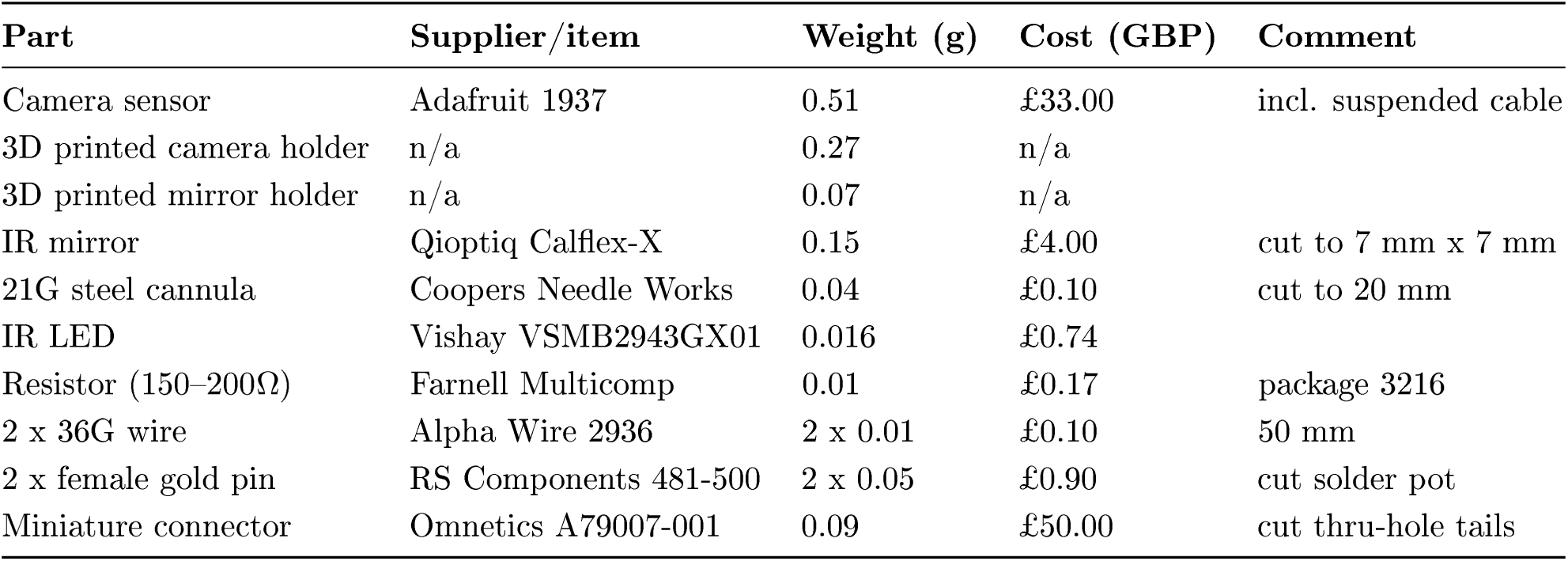
Parts required for building the miniature head-mounted camera system. Weights were measured using a calibrated micro scale (Satorius CPA225D, Goettingen, Germany). Prices for steel cannulae, mirror tiles, and wires were estimated without taking the cost of tools (e.g., glass cutter) into account. 3D printed parts were printed using a commercially available printer (Ultimaker 2+, Geldermalsen, the Netherlands) and PLA material (colorfabb, Belfeld, the Netherlands).

| Part | Supplier/item | Weight (g) | Cost (GBP) | Comment |
| --- | --- | --- | --- | --- |
| Camera sensor | Adafruit 1937 | 0.51 | £33.00 | incl. suspended cable |
| 3D printed camera holder | n/a | 0.27 | n/a |  |
| 3D printed mirror holder | n/a | 0.07 | n/a |  |
| IR mirror | Qioptiq Calflex-X | 0.15 | £4.00 | cut to 7 mm x 7 mm |
| 21G steel cannula | Coopers Needle Works | 0.04 | £0.10 | cut to 20 mm |
| IR LED | Vishay VSMB2943GX01 | 0.016 | £0.74 |  |
| Resistor (150–200Ω) | Farnell Multicomp | 0.01 | £0.17 | package 3216 |
| 2 x 36G wire | Alpha Wire 2936 | 2 x 0.01 | £0.10 | 50 mm |
| 2 x female gold pin | RS Components 481-500 | 2 x 0.05 | £0.90 | cut solder pot |
| Miniature connector | Omnetics A79007-001 | 0.09 | £50.00 | cut thru-hole tails |

## Supplemental Movies

Movie S1: **Different views with the head-mounted camera** Example views and behaviors that can be monitored using the head-mounted camera (interaction with objects, foraging, pinna movements, simultaneous monitoring of both eyes, simultaneous monitoring of eye/whisker movements and environment using two head-mounted cameras).

Movie S2: **Video image stability**. Examples demonstrating stability of the head-mounted camera system during different behaviors (locomotion, grooming, running on wheel). Shown are raw video frames (i.e. without motion correction).

Movie S3: **Continuous monitoring of behavior**. Example segment (10 minutes, playback x 25) of the data shown in Figure 5.

Movie S4: **Sound-related head and whisker movements.** An example segment of the data shown in Figure S5B,C.

Movie S5: **Eye movements in freely moving and head-fixed mice.** Eye movements measured using the head-mounted camera system (left) and for the same mouse when it was head-fixexd on a cylindrical treadmill. No stimuli or visual feedback were provided during the head-fixed recording.

Movie S6: **Prediction of eye movements.** Measured (red) and predicted (blue) eye position of a freely exploring mouse. Predictions based on a nonlinear model and head pitch/roll as shown in Figure 6D,E.

